# Rapid evolution of cone identity and retinal topography in deep-water crater lake cichlids

**DOI:** 10.64898/2026.09.16.751957

**Authors:** Monika Kłodawska, Pau Balart-García, Adrian Indermaur, Arnold Roger Bitja Nyom, Anna Nováková, Zuzana Konvičková, Oldřich Bartoš, Petr Pajer, Hana Kubová, Demian Burguera, Fanny de Busserolles, Zuzana Musilova

## Abstract

Deep-water habitats have been colonized repeatedly by fishes, driving the evolution of visual systems under extremely low-light conditions. Here, we investigated visual adaptation in cichlids from the Barombi Mbo crater lake by comparing shallow- and deep-water species using retinal single-cell transcriptomics and analyses of retinal specializations. The shallow-water species *Pungu maclareni* expressed a broader repertoire of visual opsins across distinct populations of short-(SWS), medium-(RH2), and long-wavelength-sensitive (LWS) cone photoreceptors. In contrast, the deep-water species *Myaka myaka* lacked a distinct LWS cone population and instead relied predominantly on *RH2Aα/β*-expressing, green-sensitive photoreceptors. To assess whether these deep-water RH2 cones corresponded to canonical RH2 or LWS programmes, we compared conserved photoreceptor markers. Deep-water RH2 cones retained elements of both RH2- and LWS-associated transcriptional programmes, revealing a mixed profile lacking a clear canonical cone-type signature. LWS-associated transcriptional regulators also differed between species, with *tbx2a* absent from *M. myaka* cones, consistent with the loss of LWS opsin expression, whereas *thrb* persisted in *M. myaka* RH2 cones. We further examined retinal specialization across eight species from the radiation and found that retinal topography also differed in photoreceptor distributions and ganglion cell-based acuity centers in association with habitat depth but also with trophic ecology of the species. Together, our findings show that visual adaptation in a young adaptive radiation involves coordinated changes in visual opsin expression, cone-associated transcriptional programmes, and retinal organization, highlighting how sensory systems can quickly evolve through changes at multiple biological levels in response to ecological conditions.

## Introduction

Vision is a key sensory modality underpinning survival and fitness, mediating navigation, mate choice, predator avoidance, and foraging across vertebrates [1]. In aquatic environments, visual systems are shaped by strong heterogeneity in light conditions, driven by variation in depth, clarity, turbidity, and spectral composition [2,3]. Consequently, fishes exhibit extensive visual diversification that tracks local optical environments. Adaptation to extreme habitats, such as deep or dimly lit waters, involves coordinated changes from eye morphology and retinal architecture to molecular modifications of photoreceptor genes [4,5], including opsin gene duplications, amino-acid substitutions altering spectral sensitivity, and shifts in gene expression that fine-tune visual performance under constrained light regimes [6–8].

Vertebrates possess five visual opsin classes: rhodopsin (RH1) in rods for dim-light vision, and four cone opsins (SWS1, SWS2, RH2, LWS) that contribute to spectral and colour vision [9]. Opsin expression is generally associated with five major vertebrate photoreceptor classes (i.e. rods and cones), although mismatches between opsin expression and canonical cone identity can occur [10]. Amino-acid substitutions of opsins have established distinct spectral sensitivities early in vertebrate evolution and have been repeatedly modified in response to ecological demands [11]. Teleost fishes exhibit an unusually rich opsin repertoire (median seven genes per genome [6]), exceeding that of other vertebrates and enabling adaptation across environments ranging from full-spectrum shallow-waters to spectrally restricted blue–green in deep-water habitats [12].

Cichlids are among the most diverse teleosts and are renowned for rapid adaptive radiation across gradients of depth, diet, and foraging strategy [13]. In the Barombi Mbo crater lake (Western Cameroon; Fig. 1A), an endemic flock of 11 species has diversified rapidly, exhibiting pronounced ecological and evolutionary divergence [14–16] (Fig. 1C). Molecular analyses reveal substantial visual differentiation [14]. Most shallow-water taxa express five cone opsins (*SWS2B*, *SWS2A*, *RH2Aα*, *RH2Aβ*, *LWS*), whereas deep-water species (*Konia dikume* and *Myaka myaka*) lack *LWS* expression and shift toward middle-wavelength sensitivity. In these taxa, single cones express *SWS2A*, while double cones predominantly express *RH2Aα* with lower *RH2Aβ* expression. Such depth-associated tuning likely contributed to ecological specialization within the radiation [14].

**Figure 1.**
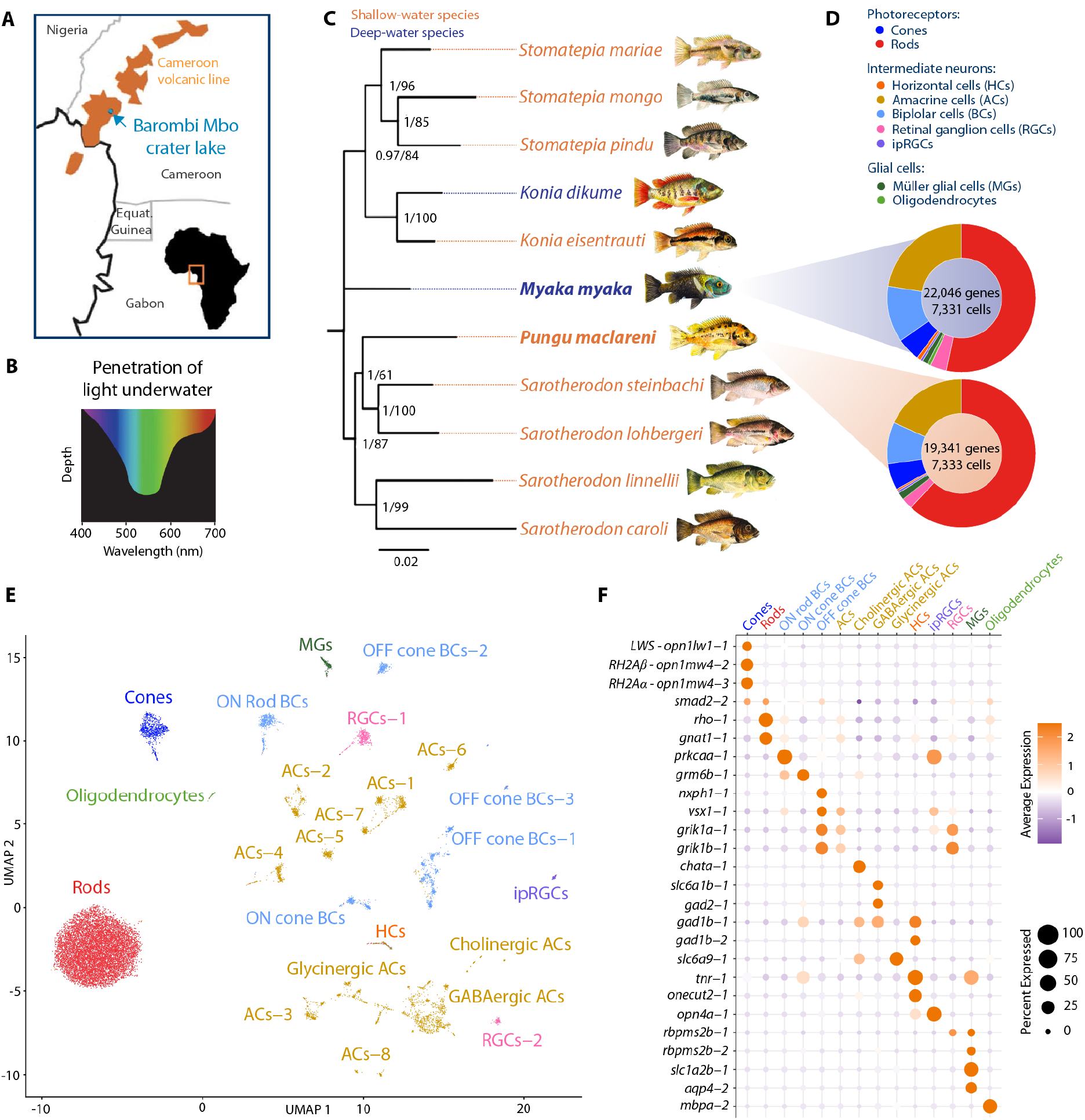
Single-cell retinal atlas of deep- and shallow-water cichlid species from the crater lake Barombi Mbo. (**A)** Geographical location of the Barombi Mbo crater lake. **(B)** Illustration of light attenuation with depth, highlighting that deeper regions of the lake receive primarily medium-wavelength (blue-green) light. **(C)** Phylogenetic tree modified from Musilova et al. (2019)[14], indicating the types of molecular (bulk RNA-seq, genome, snRNA-seq) and imaging data (quantification of retinal photoreceptors and ganglion cells) generated for each species. **(D)** Donut charts showing the proportion of major retinal cell types recovered for the two species profiled at the single-cell level: *Myaka myaka* (deep-water specialist) and *Pungu maclareni* (shallow-water specialist). **(E)** UMAP visualization of the integrated snRNA-seq dataset, with clusters annotated by retinal cell type. **(F)** Dotplot showing expression of some representative marker genes enriched in each cell type population; color gradient indicates average normalized expression levels and dot size indicates the percent of cells expressing each gene.

Although visual adaptation has been widely studied using bulk transcriptomics [17,18], single-cell approaches now resolve retinal cell-type diversity, gene regulatory programs, and photoreceptor evolution [19,20]. Nevertheless, retinal single-cell data in teleosts remain limited to a few species, including zebrafish [21], goldfish [22], grass carp, Nile tilapia, snakehead [23], and Gulf pipefish [24]. Consequently, key questions such as the allocation of opsin paralogs among photoreceptor subtypes and transcriptional specification of cone identities remain unresolved in a comparative framework.

Retinal organization evolves with ecological demands, with regions specialized for foraging, predator detection, or mate choice [25]. In fishes, spatial variation in ganglion and photoreceptor densities provides an additional level of adaptation. High ganglion cell densities (fovea or area centralis) support maximal acuity, whereas ventral or peripheral regions enhance sensitivity under dim or oblique light [3,26]. Many species exhibit multiple acuity zones or horizontal streaks reflecting behavioural specialization, and spectral sensitivity varies across the retina via opsin expression and photoreceptor density, often matching local light environments [27,28]. Together, these features illustrate how retinal architecture tunes acuity, contrast detection, and colour vision to ecological niche.

Here, we combine a chromosome-level genome assembly for *Myaka myaka* with single-cell retinal transcriptomics, spatial FISH, and stereological mapping to provide a cell-resolved view of retinal organization across the Barombi Mbo radiation. We focus on cone photoreceptor identity in two species from different depths that differ in opsin expression testing whether the deep-water middle-wavelength sensitive cones correspond to the same cone type in the shallow-water species, or alternatively, originate from the long-wavelength sensitive cones. Additionally, we clarify differential deployment of *RH2Aα* and *RH2Aβ* paralogs in double cones of shallow- and deep-water species. We further quantify photoreceptor and ganglion cell distributions in the retina to assess regional specializations in acuity and sensitivity across shallow- and deep-water species. Together, this study shows how genomic, molecular, and cellular layers of the visual system contribute to ecological divergence in this young radiation.

## Results and discussion

### A comparative atlas of retinal cell types of Barombi Mbo cichlids

To investigate retinal cell-type evolution in deep-water cichlids, we first generated a chromosome-level genome assembly for *Myaka myaka*, a deep-water specialist endemic to the crater lake Barombi Mbo (Fig. 1C; Table S1). Genome annotation, informed by retinal and multi-tissue RNA-seq and protein evidence, produced a well-supported gene set (see Methods). This resource enables species-specific single-cell RNA-seq analysis and provides a reference for comparative studies of sensory adaptation to extreme light environments.

We generated 10x Genomics single-nucleus RNA-seq data from *M. myaka* (deep-water specialist) and *Pungu maclareni* (shallow-water specialist) (Fig. 1C; Table S1), allowing comparative analysis of retinal cell types and depth-associated divergence at single-cell resolution. After quality control (removal of low-quality nuclei, ambient RNA, and doublets), we obtained datasets comprising 22,046 genes x 7,331 cells (*M. myaka*) and 19,341 genes x 7,333 cells (*P. maclareni*) (Fig. 1D). Both species showed comparable cell recovery and expressed genes, supporting robust cross-species integration (Fig. 1E).

Retinal cell types were annotated using differentially expressed genes (Fig. 1F; Table S2), guided by zebrafish markers [29] but not rigidly constrained due to phylogenetic distance. Rods were identified by strong rhodopsin expression and cones by opsin expression, whereas interneuron classes, such as amacrine and bipolar cells, were less distinct due to overlapping marker profiles. Rods dominated both datasets (53% in *M. myaka*, 62% in *P. maclareni*), followed by amacrine and bipolar cells. Cones comprised a smaller fraction (374 cells in *M. myaka*, 420 in *P. maclareni*). Oligodendrocytes were detected in *M. myaka* (55 cells) but nearly absent in *P. maclareni* (1 cell), while other retinal classes were broadly comparable between species (Fig. 1D), indicating a conserved cellular repertoire across depth gradients.

The major retinal cell types are consistent with those described in Nile tilapia snRNA-seq, which used a human-referenced annotation framework [23], and other teleost single-cell atlases [21,22,30], consistent with broad conservation of retinal cell classes across vertebrates. However, such cross-species annotation approaches may bias assignments toward conserved transcriptional programs, potentially obscuring lineage-specific features. Despite this limitation, our data show parallels with Nile tilapia, including multiple amacrine cell clusters and a dominant rod population. Whereas the tilapia dataset does not resolve amacrine subtypes, our analysis distinguishes cholinergic, GABAergic, and glycinergic AC populations, indicating a higher interneuron resolution (Fig. 1E,F). Most marker genes reported in tilapia [23] are also detected here, although paralog resolution is lacking in that study. However, several markers (e.g., *rs1a, lhx2b, nefla, sncgb, appb,* and *clu*) do not appear strictly cell-type-specific in cichlids. This is particularly evident for interneuron-associated genes (e.g., *grik1a*, *grik1b*, *gad1b*, and *tnr*), which show broader or less specific expression patterns (Fig. 1F), likely reflecting both biological divergence and differences in annotation and paralog resolution.

Overall, our data support a broadly conserved repertoire of major retinal cell classes across the two species, while revealing substantial transcriptional specialization within shared cell populations associated with their contrasting depth environments.

### Divergent cone photoreceptor repertoires in shallow- and deep-water species

To investigate how contrasting light environments are associated with cone identity, we analyzed cone photoreceptors in the shallow-water species *P. maclareni* and the deep-water species *M. myaka*. Species-specific clustering revealed clear differences in cone composition and transcriptional profiles (Fig. 2A,B; Table S3). In *P. maclareni*, we identified three major cone populations (Fig. 2A): LWS cones (long-wavelength-sensitive; also referred to as PR1 in the standardized nomenclature proposed by Baden et al. (2025) [10]; 180 cells), RH2 cones (middle-wavelength-sensitive; PR2; 172 cells), and SWS cones (short-wavelength-sensitive; PR3/4; 69 cells). In contrast, *M. myaka* contained RH2 cones (329 cells distributed across three transcriptomic clusters) and SWS cones (45 cells), but no distinct LWS cluster was detected (Fig. 2B,C). The absence of a distinct LWS population in *M. myaka* was further supported by cross-species comparison (Fig. S1; Table S4). Consistent with this difference in cone composition, *M. myaka* showed near absence of *LWS* expression and no detectable *SWS2B* expression (Fig. 2C), matching spectral constraints of deep-water light environments (Fig. 1B) [14].

**Figure 2.**
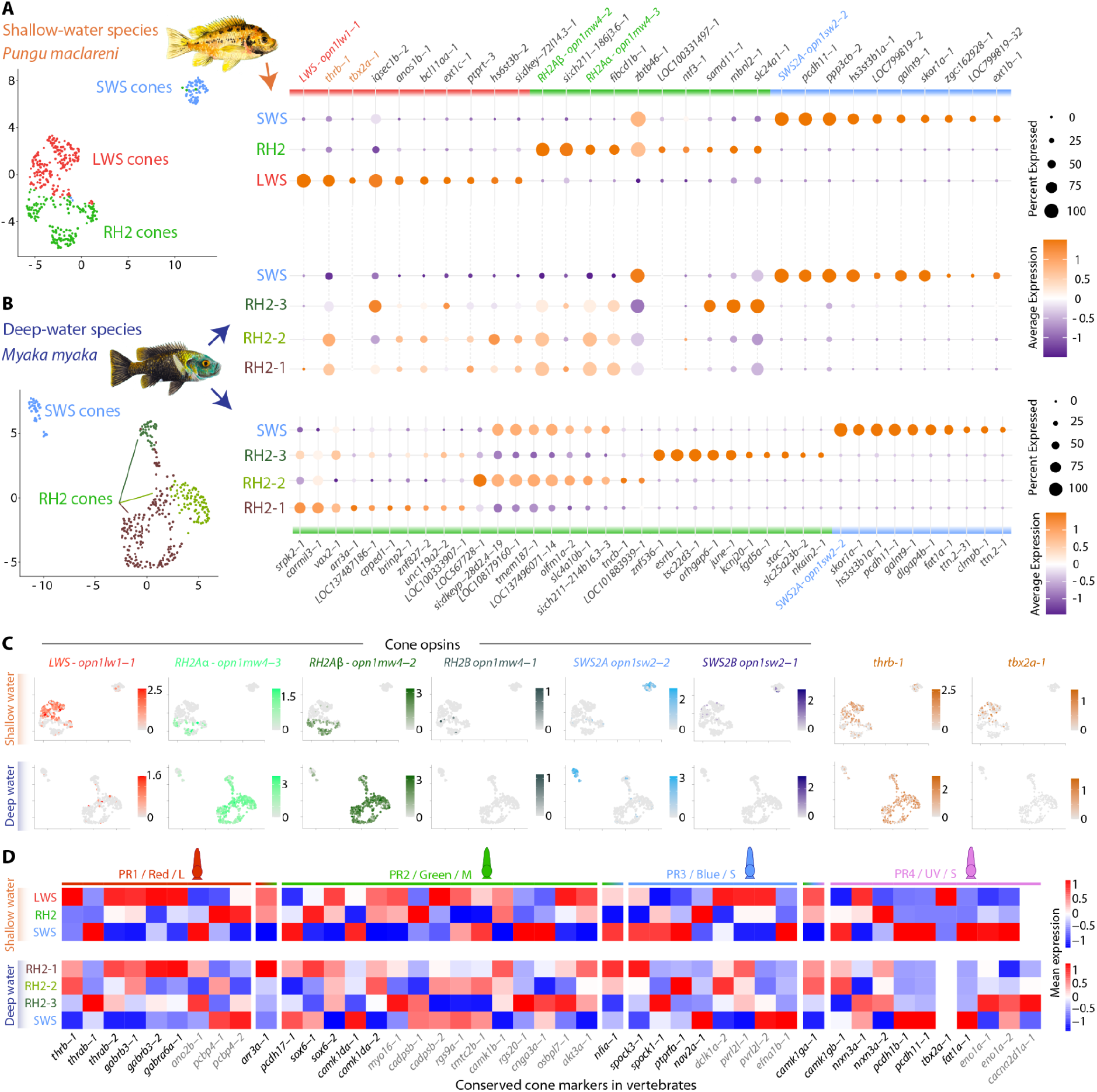
Transcriptional profiles of cone photoreceptors in deep- and shallow-water cichlids. **(A)** UMAP (left) embedding of cone photoreceptor nuclei from *Pungu maclareni* (shallow-water species), showing three canonical cone types based on opsin expression: LWS cones, RH2 cones, and SWS cones. Dotplot (right) displaying the top 10 marker gene expression for each cone cluster of *P. maclareni* (see complete list in Table S3). **(B)** UMAP (left) embedding of cone photoreceptor nuclei from *Myaka myaka* (deep-water species), showing four clusters corresponding to two cone types based on opsin expression: RH2 and SWS cones. Dotplot (right, top) projects the markers identified in *P. maclareni* onto *M. myaka* cones for direct comparison - the typical shallow-water cone identity has been remodelled. Dotplot (right, bottom) displaying the top 10 marker genes identified in *M. myaka* cone clusters (complete list in Table S3). In all dotplots, color represents average normalized expression and dot size indicates the percentage of cells expressing each gene. **(C)** Feature plots showing normalized expression of the six cone opsin genes, thyroid hormone receptor beta (*thrb-1*), and T-box transcription factor 2a (*tbx2a-1*) in both species (*P. maclareni* top, *M. myaka* bottom). Comparison across species highlights a clear separation of LWS, RH2, and SWS cones in *P. maclareni*, whereas *M. myaka* lacks a distinct LWS expression domain and shows expanded RH2-associated expression across clusters. **(D)** Heatmaps show the mean normalized expression of conserved vertebrate cone marker genes from Tommasini et al. (2025) [19] across the cone populations of *P. maclareni* and *M. myaka*, including all detected paralogs. Genes are grouped according to the four conserved photoreceptor programmes (PR1/Red/L, PR2/Green/M, PR3/Blue/S, and PR4/UV/S) defined by Baden et al. (2025) [10]. Marker genes included in the marker set by Baden et al. (2025) are labelled in black, whereas the remaining Tommasini et al. (2025) markers are labelled in grey. Expression was scaled independently for each gene across cone populations. In all plots, gene name suffixes (e.g., -1, -2) denote distinct paralogs.

The presence of both LWS and RH2 cones in the shallow-water species, but only RH2 cones in the deep-water species, raised the question of whether the latter represent an expanded RH2 cone population or instead evolved from the LWS cone lineage. To assess cone identity beyond current opsin expression, we compared the expression of conserved vertebrate photoreceptor marker sets [19] that distinguish four cone photoreceptor programmes (LWS/PR1, RH2/PR2, SWS2/PR3 and SWS1/PR4) [10]. We included the identified paralogs of these markers and compared their expression across cone populations in both species (Fig. 2D, Fig. S2). Consistent with conserved photoreceptor programmes, SWS cones in both *P. maclareni* and *M. myaka* showed strongest correspondence to PR3 and PR4 markers, with additional expression of some PR2-associated markers. *P. maclareni* LWS cones showed a clear association with PR1 markers. In contrast, RH2 cones in neither species showed a clear correspondence to a single PR programme, with markers distributed across the PR1–4 sets (Fig. 2D). These results suggest that the vertebrate PR2 marker set is less conserved or has undergone substantial reshaping in cichlids. Broader comparisons across ray-finned fish lineages, together with a fish-specific framework for cone-type homology, will ultimately be needed to determine the extent to which RH2-associated programmes are conserved across teleosts and how they relate to cone types in other vertebrates.

Beyond opsin expression, the three RH2 transcriptomic clusters in *M. myaka* showed transcriptional heterogeneity (Fig. 2B). These clusters were enriched for genes associated with phototransduction regulation (e.g. *arr3a*) [31], transcriptional control and retinal cell fate specification (e.g. *vax2*, *znf536, esrrb*) [32–34], cytoskeletal dynamics (e.g. *arhgap6*) [35], and morphogenesis (e.g. *carmil3*) [36]. However, the two *RH2Aα*- and *RH2Aβ*-expressing populations were not resolved by transcriptomic clustering but detected by FISH (see next section, Fig. 3), and the three clusters did not show distinct conserved marker signatures. We therefore interpret these clusters primarily as transcriptional subdivisions within the RH2-expressing cone population rather than as three distinct cone types. The observed differences may reflect variation in transcriptional state, regionalization, or cellular activity.

**Figure 3.**
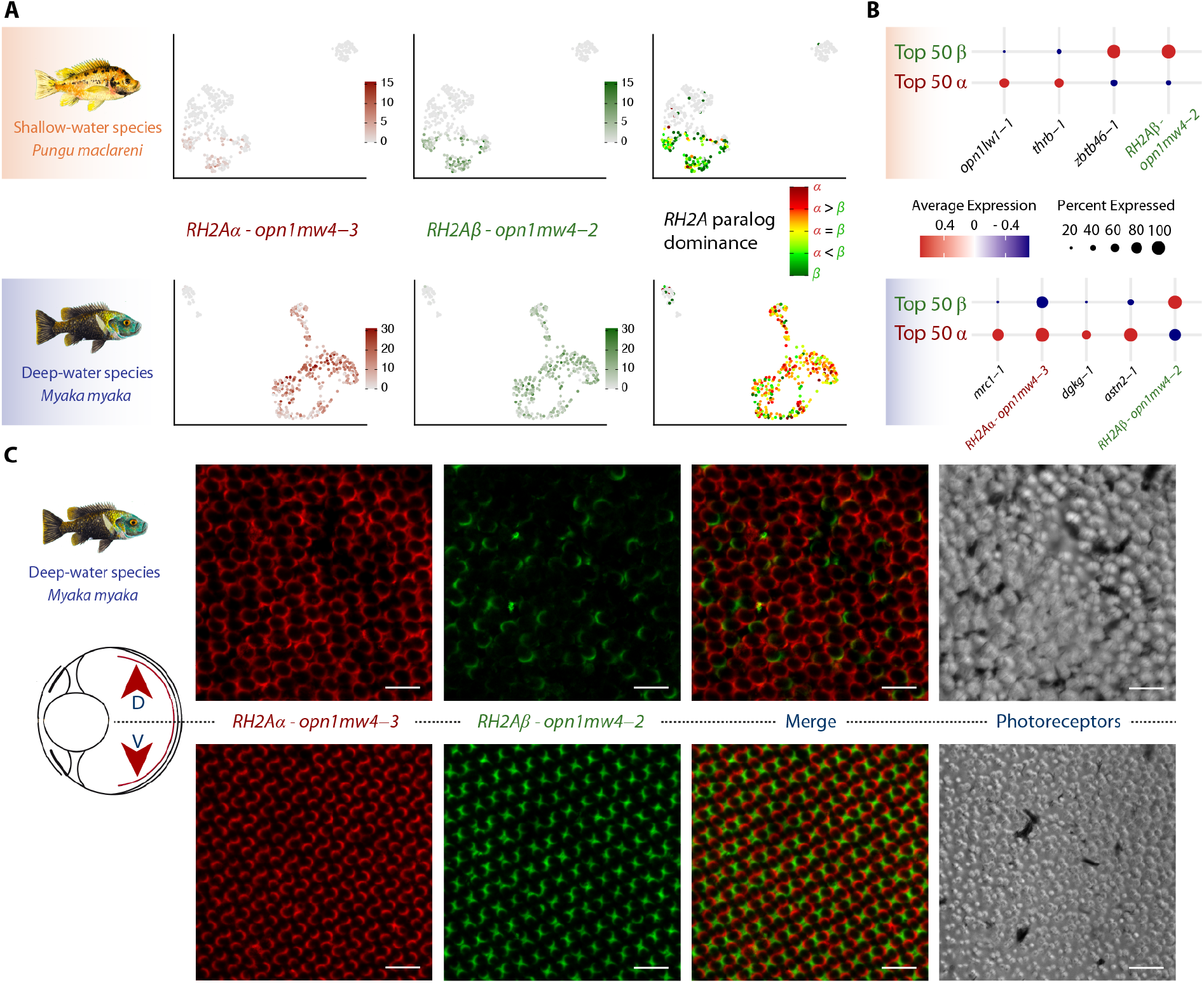
Spatial and regulatory divergence of *RH2A* opsin paralogs in cichlid retinas. (**A)** UMAP visualization of *RH2Aα* (*opn1mw4-3*) and *RH2Aβ* (*opn1mw4-2*) opsin paralogs expression in cone cells of *Pungu maclareni* and *Myaka myaka*. Individual cells are colored according to normalized RNA expression of *RH2Aα* (left, red) or *RH2Aβ* (middle, green), using the same expression scale. The right panel shows the relative *RH2A* paralog dominance, ranging from β-dominant to α-dominant, with yellow indicating approximately equal expression of the two paralogs (dominance threshold >0.2). Cells with no detectable expression of either paralog are shown in grey. **(B)** Differential gene expression analysis of the selection of 50 most α- or β-dominant *RH2A*-expressing cells in *P. maclareni* (top) and *M. myaka* (bottom). **(C)** Fluorescence in situ hybridization (FISH) using paralog-specific probes reveals spatial segregation of *RH2A* paralogs within double/twin cones in *M. myaka* (deep-water species). The ventral retina (bottom) shows a balanced ratio of *RH2Aα*- and *RH2Aβ*-expressing cones, whereas the dorsal retina (top) is biased toward *RH2Aα* expression and contains larger cells. Scale bars represent 20 μm.

Unlike *P. maclareni*, *M. myaka*, lacked a distinct *LWS*-expressing cone population, and its *RH2*-expressing cones exhibit transcriptional profiles that differed from those of both *P. maclareni* LWS and RH2 cones (Fig. 2A,B). To further characterize these differences, we performed exploratory transcriptome-wide comparisons between *P. maclareni* LWS and *M. myaka* RH2 cones (Fig. S2A), *P. maclareni* RH2 and *M. myaka* RH2 cones (Fig. S2B), and *P. maclareni* LWS and *P. maclareni* RH2 cones (Fig. S2C). These comparisons revealed substantial transcriptional differences between cone populations, with the most prominent differences involving genes outside the conserved vertebrate marker sets [19], suggesting again lack of conservation of the RH2 cone markers between cichlids and vertebrates. *M. myaka* RH2 cones differed transcriptionally from both *P. maclareni* RH2 and *P. maclareni* LWS cones, with differences involving numerous genes (Fig. S2B). Thus, *M. myaka* RH2 cones cannot be readily assigned to either *P. maclareni* cone population based on their overall transcriptional profiles.

The cone identity-associated genes showed a more nuanced pattern. Thyroid hormone receptor beta *thrb-1* was expressed in a substantial fraction of *M. myaka* RH2 cones and showed relatively similar mean expression to that observed in *P. maclareni* RH2 cones, whereas *thrab-1* was more strongly represented in *P. maclareni* RH2 cones and *thrab-2* showed similar expression between populations (Fig. S2B). Together with the vertebrate-conserved cone marker analysis, these observations indicate that *M. myaka* RH2 cones retain some components of cone-associated transcriptional programmes while lacking others. This mixed expression profile is consistent with lineage-specific modification of cone identity.

In contrast, SWS cones showed similar marker profiles across species, including *pcdh11*, *galnt9, hs3st3b1a*, together with expression of the conserved transcription factor *skor1a* [10,37] (Fig. 2A, B). These similarities indicate that SWS cones retain a more conserved molecular identity across species than was observed for RH2 cones, extending beyond differences in opsin expression.

Overall, these results show that depth-associated differences in visual systems are accompanied by shifts in opsin expression and substantial changes in cone-associated transcriptional programs. Whereas *P. maclareni* retains a broad opsin repertoire with clear cone subtype differentiation, *M. myaka* shows reduced opsin diversity and increased transcriptional heterogeneity within RH2 cones, consistent with adaptation to green-shifted deep-water light environments.

### Changes in cone-associated regulatory programs in deep-water cichlids

LWS (red-sensitive) cones are nearly absent in the deep-water specialist *M. myaka*, whereas RH2 (green-sensitive) cones dominate. To investigate the molecular basis of this shift, we examined opsin expression alongside key transcriptional regulators (Fig. 2C). The LWS-cone regulator *thrb* [38,39] is expressed in 42–88% of RH2 cones in *M. myaka* despite minimal *LWS* expression, whereas in *P. maclareni*, *thrb* is largely restricted to LWS cones (73% of this cell population). This persistence of *thrb* in *M. myaka* RH2 cones indicates retention of an LWS-associated transcriptional component despite loss of *LWS* expression, and is consistent with partial retention and remodeling of an ancestral cone regulatory programme [20,40,41].

We further identified the loss of *tbx2a* expression in *M. myaka*, a transcription factor with a well-established role in cone specification. Functional studies in teleosts have shown that loss of *tbx2a* reduces LWS cones, increases RH2 cones, and alters *thrb* expression in RH2 cones [37,42]. In zebrafish, the duplicated *tbx2a* and *tbx2b* paralogs have partially diverged roles in cone identity: *tbx2a* is enriched in LWS cones and contributes to their identity, whereas *tbx2b* is enriched in SWS cones and contributes to SWS-cone identity; both paralogs are also independently required for UV-cone generation [37]. Consistent with these paralog-specific roles, single-nucleus RNA-seq revealed a striking absence of *tbx2a* in *M. myaka* cones, in contrast to its strong and LWS-specific expression in *P. maclareni* (Fig. 2C, Fig. S3A). In contrast, *tbx2b* was conserved between the species, being enriched in SWS cones and detected at lower levels in RH2 and LWS cones. Other transcriptional differences included higher *rx1-1* expression in *P. maclareni* LWS cones and higher *rx1-2* and *neurod1-1* expression in *M. myaka* RH2 cones, whereas most other retinal transcription factors showed low and relatively uniform expression across cone types (Fig. S3A). Together, these differences point to specific remodelling of cone-associated transcriptional programmes accompanying the loss of LWS expression in *M. myaka*.

Notably, *tbx2a* was included among the PR4 markers in Tommasini et al. (2025) [19]. We retained *tbx2a* in this marker set for the direct comparison with their framework, but its expression in our data is strongly associated with *P. maclareni* LWS cones rather than SWS cones (Fig. 2D). This illustrates that individual conserved markers may not retain identical cone-type associations across vertebrate lineages and should therefore be interpreted in the context of the broader marker programme. In *M. myaka*, the absence of *tbx2a* together with persistence of *thrb* provides an example of the selective retention and loss of cone-associated regulatory components during the evolutionary shift toward RH2 expression.

To assess whether these changes were accompanied by broader alterations in regulatory-network activity, we performed a gene regulatory network (GRN) analysis across LWS and RH2 cone populations in *M. myaka* and *P. maclareni*. Only three regulons were robustly recovered due to cross-species motif mapping constraints, including *Tbx2* (Fig. S3B). *Tbx2* regulon activity was highest in *P. maclareni* LWS cones, but did not differ significantly from *M. myaka* RH2 cones (i.e., BH-adjusted pairwise Wilcoxon test, P = 0.876), whereas activity was significantly higher in *P. maclareni* LWS *than* RH2 cones (P = 0.036). *M. myaka* RH2 also showed higher *Tbx2* regulon activity than *P. maclareni* RH2 (P = 0.019). Thus, despite the absence of detectable *tbx2a* expression in *M. myaka*, the inferred *Tbx2* regulon activity in its RH2 cones was comparable to that of LWS cones. Other regulons (*Foxp1*, *Sox2*) showed uniform activity across species and cones populations. These results provide limited evidence for differences in inferred regulon activity and do not support broad remodeling of the regulatory network, although the small number of robustly recovered regulons precludes a comprehensive assessment of regulatory-network changes.

Comparative genomic analyses indicate that *tbx2a* silencing in *M. myaka* is not due to gene decay; the coding sequence remains intact and highly conserved relative to *P. maclareni* as well as *Metriaclima zebra* (i.e., a shallow-water cichlid from the Malawi lake) [42]. Instead, *M. myaka* harbors lineage-specific insertions in upstream regulatory regions. Motif mapping shows that a proximal insertion (+16 bp at –612 bp) introduces additional *klf9* binding sites, enriched in *M. myaka* (Fig. S3C; Table S5), whereas more distal insertions generate additional binding sites for other factors, though these are fewer and likely less relevant to cone regulation. *klf9* is a context-dependent transcriptional repressor in vertebrates [43,44], though its role in cone photoreceptors remains untested. These observations are consistent with a potential cis-regulatory contribution to the reduced *tbx2a* expression in *M. myaka*, but functional experiments will be required to establish causality.

Despite *tbx2a* silencing, *TBX2*-family binding sites are retained in the regulatory regions of the *LWS* and *RH2Aβ* opsins in *M. myaka*, whereas *RH2Aα* lacks such motifs, consistent with observations in Malawi cichlids [42] and suggesting paralog-specific regulatory sensitivity. This preservation of downstream regulatory architecture, alongside upstream *tbx2a* silencing, indicates that altered opsin expression primarily reflects regulatory changes rather than mutations at opsin loci. Functional studies showed that *tbx2a* loss reduces LWS cones, increases RH2 cones, and alters *thrb* expression in RH2 cones [37,42]. Collectively, these findings identify specific changes in cone-associated regulatory components accompanying the reduced opsin repertoire of this deep-water species.

### Spatial segregation of *RH2A* opsin paralogs across the retina

Single-nucleus RNA-seq revealed distinct patterns of relative *RH2Aα* and *RH2Aβ* paralog expression among RH2 cones of *M. myaka* and *P. maclareni* (Fig. 3A). Using a threshold (D>0.2) to define paralog-dominant cells, *M. myaka* showed a relatively balanced distribution, with 47.4% of RH2 cones α-dominant, 23.1% β-dominant, and 29.4% showing approximately balanced expression of both paralogs. In contrast, *P. maclareni* showed a pronounced bias toward *RH2Aβ*, with 78.3% of cells β-dominant, compared with only 7.2% α-dominant and 14.5% balanced. Thus, *RH2A* paralog usage differs markedly between species, with *M. myaka* showing substantial representation of both α- and β-dominant cells and a relatively large balanced population, whereas *P. maclareni* is strongly biased toward *RH2Aβ* expression. Given the high sequence similarity between paralogs (∼98%) we considered potential read mapping ambiguity, however after thorough check of the 3’UTR region, the paralog resolution in the snRNA-seq data should be sufficient (Fig. S4).

To test whether paralog dominance corresponded to broader transcriptional differences, we performed differential expression analysis on the 50 cells with the strongest relative expression of each paralog (Fig. 3B; Table S6). In *M. myaka*, only three genes (*mrc1*, *dgkg*, and *astn2*) were enriched in *α*-dominant cells, and none in *β*-dominant cells, indicating that *RH2A* paralog usage is largely uncoupled from broader transcriptional differences among RH2 cones. In *P. maclareni, α*-dominant cones were enriched for *opn1lw1* and *thrb*, whereas *β*-dominant cones showed enrichment of *zbtb46* alongside *RH2Aβ* (Fig. 3B; Table S6). This pattern suggests partial co-expression of *LWS* and *RH2Aα* within a subset of cones rather than discrete cone types, consistent with the absence of clear clustering. Thus, in *P. maclareni*, *RH2A* paralog expression remains embedded within a broader cone identity, whereas in *M. myaka* this regulatory coupling appears reduced.

To further resolve spatial organization of the two *RH2A* paralogs, we performed fluorescence in situ hybridization (FISH) with paralog-specific probes. In *M. myaka*, double cones showed consistent asymmetry, with one member predominantly expressing *RH2Aα* and the other *RH2Aβ*, particularly in the ventral retina, whereas the dorsal retina was dominated by *RH2Aα-*expressing cones (Fig. 3C). The FISH results therefore support the single-cell RNA-seq evidence for asymmetric paralog usage, while additionally revealing its organization within individual double/twin cones and across retinal regions. The subset of approximately balanced cells detected by snRNA-seq may reflect genuine co-expression, but differences between the two approaches could also arise because snRNA-seq primarily samples nuclear transcripts, whereas FISH detects cellular mRNA.The enrichment of *RH2Aβ*-expressing cones in the ventral retina suggests functional regionalization. In teleosts, the ventral retina samples downwelling light [45], and *M. myaka*, a pelagic zooplanktivore [46] with an upward-facing mouth, likely relies on detecting prey against brighter backgrounds. Spatial partitioning of *RH2A* paralog expression may therefore fine-tune spectral sensitivity within the green-shifted deep-water light. This interpretation is consistent with regional variation in photoreceptor and ganglion cell densities in *M. myaka* (see below), indicating coordinated spatial adaptation of retinal architecture.

### Seasonal modulation of photoreceptor investment

*Myaka myaka* is a seasonally migratory species that resides in deep waters (∼20 m) of the lake Barombi Mbo for most of the year but moves into shallow littoral zones during the rainy season to reproduce [14,46]. During this period, males defend territories while females roam among them to spawn. To assess seasonal plasticity in visual gene expression, we generated bulk RNA-seq data and compared opsin profiles of individuals collected in the dry (deep-water) and rainy (shallow-water) seasons.

Dry-season individuals, corresponding to deep-water residency, showed higher relative rod rhodopsin expression compared to cone opsins, consistent with increased reliance on scotopic vision under low-light. In contrast, rainy-season individuals exhibited a shift toward cone-dominated expression, with an increase in single-cone opsins relative to double-cone opsins (Table 1). A similar seasonal increase in single-cone opsin representation, but not rod-cone balance, has been reported in the crater lake radiation of *Coptodon* cichlids from Lake Bermin [18]. These results indicate pronounced seasonal plasticity in photoreceptor investment, likely reflecting reversible adjustments to changing light environments.

**Table 1.** Seasonality shifts in the proportional opsin expressions in *Myaka myaka*. Mean and SD proportional expression (%) of retinal opsins in individuals sampled during the rainy season (shallow water) and dry season (deep water). Rod and cone proportions are expressed relative to total opsin expression. Single- and double-cone proportions are expressed relative to total cone opsins. Individual opsin classes are expressed relative to their respective cone type. P-values indicate seasonal comparisons (dry vs. rainy). ***P < 0.001.

|  |  | Rainy season<br>(shallow water) |  | Dry season<br>(deep water) |  |  |
| --- | --- | --- | --- | --- | --- | --- |
|  |  | Mean | SD | Mean | SD | dry vs. rainy |
| **Rod and cone proportions from total opsins (%):** | rods | 86.3 | 5.0 | 98.1 | 2.0 | *** |
|  | cones | 13.7 | 5.0 | 1.9 | 2.0 |  |
| **Single- and double cones from all cone opsins (%):** | single | 24.3 | 6.3 | 13.3 | 5.9 | *** |
|  | double | 75.7 | 6.3 | 86.7 | 5.9 |  |
| **Double-cone opsins** | LWS | 0.4 | 0.6 | 0.1 | 0.1 |  |
| | RH2Aα | 78.3 | 8.4 | 79.1 | 8.4 | |
| | RH2Aβ | 21.3 | 8.2 | 20.1 | 8.2 | |
|  | RH2B | 0.1 | 0.1 | 0.7 | 0.6 |  |
| **Single-cone opsins** | SWS1 | 0.0 | 0.0 | 1.2 | 2.1 |  |
|  | SWS2A | 97.9 | 1.1 | 94.5 | 4.1 |  |
|  | SWS2B | 2.1 | 1.1 | 4.3 | 4.0 |  |

Despite these shifts, *RH2Aα* and *RH2Aβ* expression remained stable across seasons, and *SWS2B* and *LWS* opsins remained minimally expressed in both conditions. Thus, while *M. myaka* exhibits seasonal modulation of rod–cone balance and cone subtype composition, its opsin repertoire and *RH2A* paralog usage are largely invariant. This combination of plastic and stable traits suggests that seasonal habitat transitions are accommodated primarily through quantitative changes in photoreceptor deployment rather than changes in spectral tuning.

### Visual sensitivity vs. acuity: the topographic distribution of photoreceptor and ganglion cells aligns with depth specializations

Distinct visual tasks drive regional retinal specialization [47]. To assess how retinal architecture varies with habitat depth, we reconstructed the topographic distributions of cone photoreceptors and retinal ganglion cells across Barombi Mbo cichlids spanning the shallow–deep gradient. Photoreceptor topography was analysed in two shallow- and two deep-water species, whereas ganglion cell distributions were quantified across eight species (Fig. 4).

**Figure 4.**
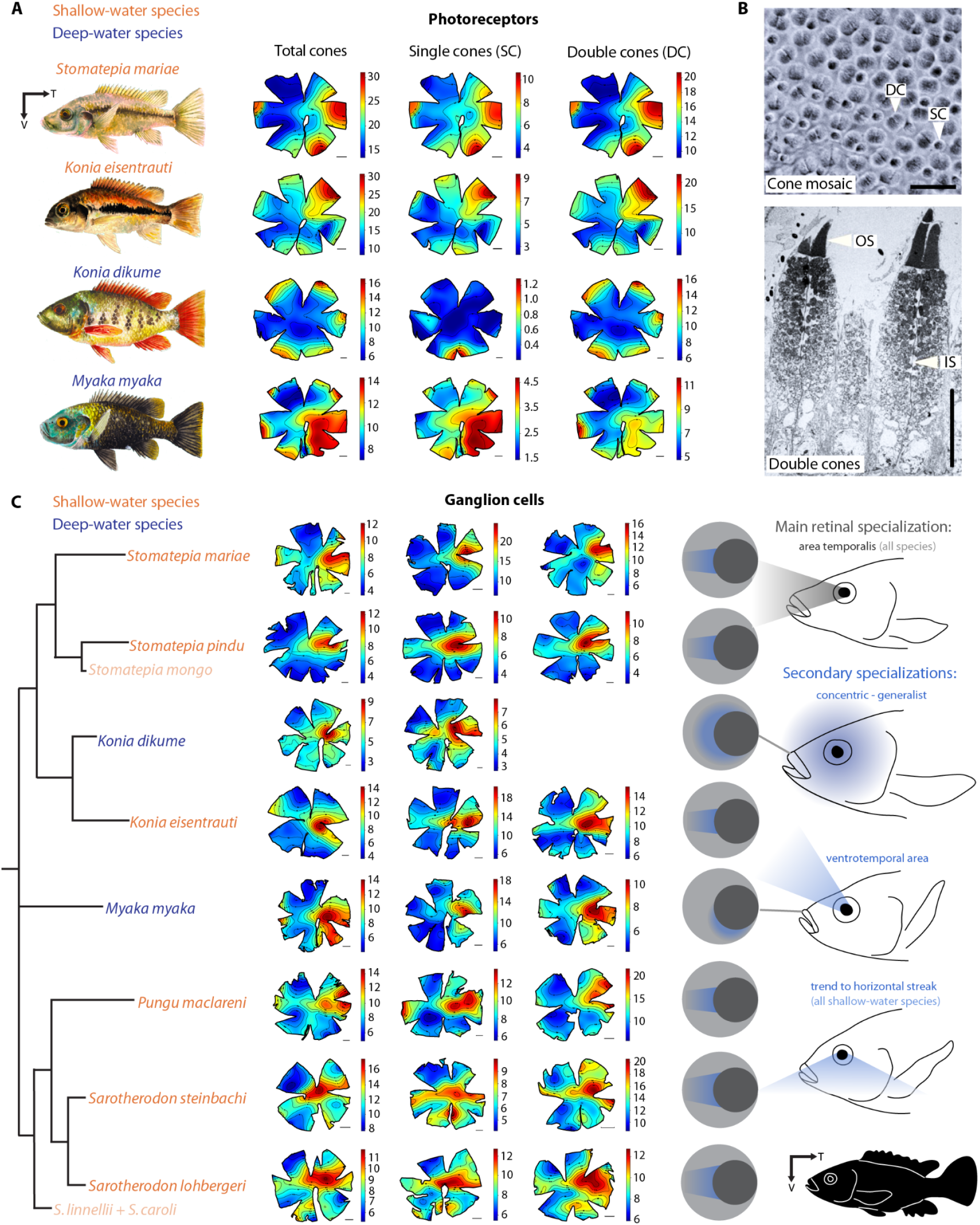
Retinal topographic maps of Barombi Mbo cichlids. **(A)** Photoreceptor cell density in shallow-water and deep-water species. Only one individual per species is shown, additional individuals are included in figure S3. **(B)** Retinal photoreceptors in cichlids: whole mount image of mosaic of single and double cones in *Stomatepia mariae* (top) and transmission electron micrograph (TEM) of two double cones in *Konia eisentrauti* (bottom). DC = double cone, SC = single cone, OS = outer segment, IS = inner segment. **(C)** Ganglion cell density maps highlighting the areas of highest visual acuity in eight cichlid species, showing 2 or 3 replicates per species. All Barombi cichlids share the main retinal specialization (highest acuity) in the temporal area, but they differ in the secondary specializations. All shallow-water species show a trend towards a horizontal streak (more pronounced in the two *Sarotherodon* species), whereas the two deep-water species show different secondary specialization: *Konia dikume* has a concentric density pattern usually found in visual generalists, whereas the area of *Myaka myaka* tends to extend ventrally – corresponding possibly to the upward oriented mouth shape and, hence, its most common feeding direction in the deep pelagic zone. T= Temporal, V = Ventral. Scale bars in (a) and (c) represent 1 µm. Scale bar in (b) represents 20 µm (top) and 10,000 nm (bottom).

Across species, double cones dominated the photoreceptor class (∼78% of all cones; Table 2), consistent with transcriptomic data. All species except *K. dikume* exhibited an arch specialisation, with increased cell density along the dorsal-temporal-ventral area and species-specific peak positions, consistent with enhanced forward-directed vision (Fig. 4A; Fig. S5). Such curved regions of elevated density likely enhance spatial resolution across defined portions of the visual field and resemble photoreceptor specializations described in lanternfishes [47]. In contrast, the obligate deep-water specialist *K. dikume* showed broadly elevated double-cone densities across much of the retinal periphery, consistent with continuous ocular growth and peripheral photoreceptor addition in teleosts [48]. This pattern likely reflects a sensitivity-biased strategy in dim-light environments, where larger cones increase photon capture, balancing size and number [47,49].

**Table 2.** Photoreceptor and ganglion cell counts in the retina of Barombi Mbo cichlids. (see Table S7 for additional details). SRP = spatial resolving power.

| Photoreceptor cells: |  |  |  |  |  |  |
| --- | --- | --- | --- | --- | --- | --- |
| Species | Ecology | ID | Estimated total number | double cones | single cones | AVG single cones |
| *Konia eisentrauti* | shallow water | 464 | 599,885 | 68% | 32% | 30.70% |
|  |  | 644 | 1,022,363 | 69% | 31% |  |
|  |  | M199 | 555,883 | 71% | 29% |  |
| *Stomatepia mariae* | shallow water | H079 | 690,081 | 68% | 32% | 23.70% |
|  |  | A1 | 831,797 | 75% | 25% |  |
|  |  | A2 | 1,266,324 | 86% | 14% |  |
| *Konia dikume* | deep water | 490 | 723,278 | 94% | 6% | 4% |
|  |  | 522 | 682,776 | 99% | 1% |  |
|  |  | 630 | 667,950 | 95% | 5% |  |
| *Myaka myaka* | deep water | 596 | 562,651 | 68% | 32% | 31.30% |
|  |  | M1R | 765,772 | 69% | 31% |  |
|  |  | M3L | 697,388 | 69% | 31% |  |
| Retinal ganglion cells: |  |  |  |  |  |  |
| Species | Ecology | ID | Estimated total number | Peak density (cells/mm2) | SRP (cpd) of visual arc | mean SRP |
| *Konia eisentrauti* | shallow water | 512 | 537,440 | 33,594 | 5.92 | 5.39 |
|  |  | 475 | 501,368 | 22,813 | 4.82 |  |
|  |  | H069 | 630,251 | 19,063 | 5.44 |  |
| *Pungu maclareni* | shallow water | M208 | 432,544 | 40,313 | 5.31 | 5.35 |
|  |  | P3R | 664,140 | 17,813 | 5.06 |  |
|  |  | M089 | 730,496 | 23,438 | 5.68 |  |
| *Sarotherodon lohbergeri* | shallow water | M177 | 367,632 | 18,281 | 4.06 | 3.96 |
|  |  | S2L | 400,105 | 15,469 | 3.7 |  |
|  |  | M167 | 517,369 | 15,156 | 4.12 |  |
| *Sarotherodon steinbachi* | shallow water | M097 | 255,220 | 28,750 | 3.65 | 3.81 |
|  |  | 477 | 428,331 | 14,219 | 4.06 |  |
|  |  | M170 | 371,252 | 22,188 | 3.71 |  |
| *Stomatepia mariae* | shallow water | M095 | 666,304 | 20,000 | 5.44 | 5.58 |
|  |  | 585 | 841,825 | 18,438 | 6.24 |  |
|  |  | M093 | 408,166 | 35,625 | 5.06 |  |
| *Stomatepia pindu* | shallow water | M084 | 547,366 | 20,469 | 5.96 | 5.1 |
|  |  | H019 | 481,584 | 15,156 | 4.64 |  |
|  |  | M264 | 433,147 | 16,406 | 4.7 |  |
| *Konia dikume* | deep water | 552 | 665,864 | 13,438 | 5.66 | 5.7 |
|  |  | 626 | 676,306 | 13,906 | 5.74 |  |
| *Myaka myaka* | deep water | 615 | 503,733 | 12,716 | 4.15 | 4.86 |
|  |  | 579 | 480,997 | 21,235 | 4.75 |  |
|  |  | 523 | 440,151 | 28,889 | 5.68 |  |

Single cones in *K. dikume* were strongly concentrated in the ventral retina and largely absent elsewhere (Fig. 4A), a pattern not observed in other Barombi species. Comparable spatial heterogeneity has been reported in other fishes, including deep-sea taxa with localized cone patches in otherwise rod-dominated retinas [6,50], analogous to the pattern observed in *K. dikume*. In both cases, the ventral retina is oriented to receive downwelling light from the surface [51], likely enhancing sensitivity under threshold light conditions. Consistent with this, the double-cone to single-cone (DC/SC) ratio varied markedly among species: *K. dikume* showed an extreme bias towards double cones (DC/SC ≈ 24:1), whereas all other taxa, including the seasonally deep-water species *Myaka myaka*, exhibited rations of ∼3–4:1 (Table 2). Despite sharing an identical opsin expression repertoire (*SWS2A* in single cones; *RH2Aα*/*β* in double cones) [14–16], *M. myaka* and *K. dikume* differ strongly in photoreceptor topography. This decoupling of spectral tuning from retinal organization indicates that similar chromatic sensitivities can arise through distinct spatial configurations, likely reflecting differences in depth use, ecology, and visual behaviour.

Ganglion cell topography was examined in six shallow-water species (*Stomatepia mariae*, *S. pindu*, *Konia eisentrauti*, *Pungu maclareni*, *Sarotherodon steinbachi*, *S. lohbergeri*) and two deep-water species (*Konia dikume*, *Myaka myaka*). All species possessed a pronounced area temporalis as the primary region of highest visual acuity (Fig. 4C). Secondary specializations differed among ecological groups. Shallow-water species exhibited a weak horizontal streak, more pronounced in *Sarotherodon*, whereas deep-water species displayed distinct secondary specializations, including concentric or ventrally extended regions of increased acuity (Fig. 4C). The conserved temporal specialization, coupled with divergence in secondary specializations likely reflects the young evolutionary age of the radiation.

In teleosts, retinal topography is closely linked to trophic ecology, with temporal specializations commonly associated with active predation and forward-directed vision [28,45,52,53], and representing a predominant arrangement in reef fishes [28,54]. All Barombi species retain this feature, likely inherited from a common ancestor, as the closely related Nile tilapia (*Oreochromis niloticus*) shows similar organization [55].

The horizontal streak observed in shallow-water species is associated with wide-angle vision without eye movements [45,47], and facilitates scanning the horizon, as seen in open-water swimmers. Consistent with this, *S. steinbachi* forms surface-foraging schools, whereas *S. lohbergeri* schools roam the water column [46].

Deep-water species lack a horizontal streak and instead exhibit distinct secondary specializations. *K. dikume*, a strict deep-water resident, shows a concentric pattern of retinal acuity consistent with a generalized visual strategy. By contrast, *M. myaka*, a seasonally deep-water species, exhibits a ventrally extended specialization likely enhancing detection of objects against downwelling light, consistent with its zooplanktivorous, upward-directed feeding behavior (Fig. 4C). Similar links between retinal topography and feeding ecology have been described in lanternfishes [25], holocentrids [56], and cardinalfishes [57], although the mechanisms underlying regional specialization remain poorly resolved [3,56,58]. Interestingly, in *K. dikume*, photoreceptor (see above) and ganglion cell topographies were not spatially matched (Fig. 4A,C), unlike the pattern typical of teleosts, although similar mismatches occur in nocturnal holocentrids [56].

To further assess adaptive divergence, we quantified visual acuity from ganglion cell densities. In deep-water environments, *K. dikume* experiences strong selection for sensitivity, favoring increased photoreceptor convergence at the expense of ganglion cell density. Despite reduced ganglion cell densities (Table 2), it exhibited the highest spatial resolving power (SRP; 5.70 cycles deg⁻¹, hereafter cpd), supported by larger eyes and pupil diameters that increase photon capture (table S7). The visually oriented predator *Stomatepia mariae* showed similar high SRP (5.58 cpd), whereas omnivorous and herbivorous *Sarotherodon* displayed lower values (3.81–3.96 cpd; Table 2).

These patterns indicate that retinal architecture and ocular scaling have diverged in accordance with ecological niche [45], with high spatial resolution in predators (*Stomatepia*) and deep-water specialists (*K. dikume*), and lower acuity in taxa less reliant on visually mediated prey detection (*Sarotherodon*). SRP values fall within the range reported for other cichlids (e.g., *Metriaclima zebra*, ∼4.5 cpd; Victorian cichlids, 3.3–3.9 cpd) [3,59] and are comparable to other reef teleosts (e.g., blennies, sandperches), although some taxa (e.g., emperors, tuskfishes) exhibit substantially higher acuity [52].

Overall, retinal organization in Barombi Mbo cichlids closely reflects ecological and visual demands. Shallow-water species exhibit localized photoreceptor and ganglion cell specializations supporting forward-directed, high-acuity vision with a secondary weak horizontal streak, whereas deep-water species lack this feature and instead show broader photoreceptor distributions and, in *K. dikume*, extreme double-cone dominance, consistent with a sensitivity-biased strategy. Together with species-specific variation in spatial resolving power, these patterns demonstrate how retinal architecture and ocular scaling have diverged across the radiation to match habitat depth and foraging ecology.

Our comparative analysis of deep- and shallow-water cichlids shows that adaptation to contrasting light environments involves coordinated changes in opsin expression, cone-associated transcriptional programmes, and retinal spatial organization. In the deep-water specialist *Myaka myaka*, spectral tuning involves loss of *LWS* expression, altered *RH2A* paralog usage, and selective changes in cone-associated transcriptional components, including loss of *tbx2a* expression alongside persistence of *thrb* in RH2-expressing cones. Conserved cone-marker analysis further reveals a mixed cone-associated transcriptional signature in *M. myaka* RH2 cones, without supporting a simple canonical cone-type assignment. Together, our findings show that visual adaptation to deep-water habitats involves coordinated but not strictly coupled modifications across multiple levels of retinal organization, providing a framework for understanding ecological specialization along light gradients.

## Materials and Methods

### Animal collection and sample preservation

Samples of adult individuals were collected at the crater lake Barombi Mbo (Cameroon) during both dry (November–March) and rainy (April–October) seasons, under the following research permits: 0000047,49/MINRESI/B00/C00/C10/nye, 0000116,117/MINRESI/B00/C00/C10/C14, 000002-3/MINRESI/B00/C00/C10/C11, 0000032,4850/MINRESI/B00/C00/C10/C12). To minimize potential circadian effects on gene expression, most samples were collected within a consistent time window between 7:00 and 13:00. Fish were captured using selective gill netting while snorkelling. Following euthanasia (anaesthetic overdose or percussive stunning), eyeballs were dissected from freshly euthanized individuals. Eyes were opened with a dorsal incision through the pupil to preserve orientation and fixed with 2% PFA and 2.5% glutaraldehyde in PBS for histological analyses, and with 4% PFA in PBS which was changed for 1g/L NaN₃ in PBS for storage for photoreceptors and ganglion cells examination. For transcriptomic analyses, whole eyeballs were preserved in RNAlater™ (Invitrogen™). Some *Myaka myaka* and *Pungu maclareni* individuals were kept alive and transported to the laboratory aquarium for downstream genomic, single-nucleus RNA-seq, and fluorescent in-situ hybridization analyses.

### Genome sequencing and assembly

High-molecular weight genomic DNA from *Myaka myaka* was isolated from flash-frozen fin tissue. Tissue was pulverized in liquid nitrogen using a pre-cooled mortar and pestle, and the resulting powder was lysed overnight at 55°C in lysis buffer (250 mM EDTA pH 8.5, 10 mM CaCl₂, 1% SDS, 200 µg/ml proteinase K). Following RNase A treatment and phenol/chloroform extraction, DNA was precipitated with ethanol and recovered either by spooling onto a glass microstirrer or, when necessary, by centrifugation. The pellet was dissolved in TE buffer (10 mM Tris-HCl pH 8.5, 1 mM EDTA, 10 mM NaN₃), and DNA concentration was quantified using a Qubit fluorometer. Long-read sequencing was performed on a GridIONx5 platform (Oxford Nanopore Technologies) using R9.4.1 flow cells and chemistry. Basecalling was performed with Guppy v3.2.10 using the high-accuracy (HAC) model. The resulting dataset comprised approximately 45.3 Gb of data (∼45× estimated genome coverage), with read N50 and N90 lengths of 7,024 bp and 2,579 bp, respectively. De novo genome assembly was carried out with Flye v2.9.1 [60] in nano-raw mode, using an expected genome size of 1.0 Gb and a minimum overlap length of 5,000 bp. This step produced a primary long-read assembly used for subsequent polishing. The initial Flye assembly was polished with Medaka v1.11.3 to correct systematic ONT basecalling errors, and a consensus sequence was generated to improve base-level accuracy prior to short-read polishing. Additional polishing was performed using Illumina Hi-C reads, which provided high-depth short-read coverage (∼90× mean coverage). Polishing was carried out with four iterative rounds of Pilon v1.24, focusing exclusively on base-level corrections (--fix bases). Next, paired-end Hi-C reads were aligned using BWA-MEM [61], and only mapped, non-duplicate read pairs were retained for scaffolding. Chromosome-scale scaffolding was achieved using YaHS, which leverages Hi-C contact information to order and orient contigs. Manual inspection and correction of scaffolds were performed using Juicebox Assembly Tools (JBAT) [62]. Scaffolds shorter than 15 kb were removed. The final assembly spans 0.92 Gb across 865 scaffolds, with an N50 of 37.4Mb, and shows a high completeness based on BUSCO Actinopterygii odb10 database (98.9% complete [single-copy: 98.1%, duplicated: 0.8%], 0.3% fragmented, 0.8% missing; n = 3640).

### Genome annotation

The genome assembly was masked using a repeat library generated with RepeatModeler v.1.0.11 [63] and subsequently applied with RepeatMasker v.4.0.7 (www.repeatmasker.org). Total RNA from different adult tissues (retina, gills, and olfactory epithelium samples) was extracted using the RNeasy Mini Kit (Qiagen). Libraries were prepared and sequenced by Novogene on an Illumina platform, producing short paired-end reads (150 bp × 2). The extracted RNA concentration and integrity were verified using a 2100 Bioanalyzer (Agilent) and Qubit Fluorometer (Thermo Fisher Scientific). Additional information is listed in Table S1. We annotated the genome of *Myaka myaka* by integrating RNA-seq and protein evidence. RNA-seq was used for annotation after passing a quality check using Bioanalyzer (RIN ≥ 7). Raw reads were processed with fastp v.0.22.0 [64] to remove low-quality sequences and adapters, retaining reads longer than 50 bp. Filtered reads were aligned to the reference genome using HISAT2 v.5.3.1 [65], applying options to report transcriptome-optimized alignments (--dta) and suppress unpaired or discordant alignments (--no-mixed, --no-discordant). Then we used Samtools v.1.14[65] for converting SAM outputs to BAM, filtering properly aligned reads (i.e., options -F3844 and -q 60), and concatenating all files into a single sorted BAM file for downstream analysis. A manual inspection step with IGV was performed to discard datasets with low mapping rate and or bad transcript reconstruction. Transcript assembly was performed using StringTie v2.2.1 [66] on the merged BAM, initially with default parameters; however, transcript fusion issues led to the adoption of more stringent settings (i.e., -c 1.5, -f 0.1, -m 300, -j 3, -a 20), generating a conservative GTF annotation suitable for subsequent steps. Alternatively, protein-coding gene prediction was performed using BRAKER3 [67], integrating RNA-seq evidence and protein homology using *Oreochromis niloticus* proteome (GCF_001858045.2) to aid predicting gene models. We used AGAT v1.4.1 (https://github.com/NBISweden/AGAT) to merge StringTie and BRAKER3 annotations and to process GFF and GTF files. Coding sequences were predicted with TransDecoder v.5.7.0 (https://github.com/TransDecoder/TransDecoder), integrating BLAST v.2.10.0 [68] (i.e., using a database based on *O. niloticus* proteome) and HMMER v.3.3.2 [69] (i.e., using the Pfam-A.hmm database). The TransDecoder output was annotated using blastp with evalue 1e-10 and selecting a single hit per sequence based on the bitscore with primarily (i) the NCBI Zebrafish proteome (i.e., assembly GRCz11), and complemented by additional hits found with (ii) the UniProt Zebrafish proteome (UP000000437). The mitochondrial genes were annotated using the MITOfish tool [70]. We employed AGAT to resolve overlapping genes and standardize GTF formatting, followed by custom Perl scripts to correct transcript fusions and to handle inconsistencies on strand assignments. To ensure compatibility with downstream single-cell analyses, duplicated gene names were uniquely suffixed and a transcript-to-gene table was generated. Finally, CDS sequences corresponding to annotated transcripts were filtered to create a FASTA file suitable for single-cell analyses. This pipeline allows for robust integration of transcriptomic and proteomic evidence, producing a curated and non-redundant gene annotation suitable for functional genomics and single-cell approaches.

### Nuclei isolation, sorting and sequencing

Retinae of adult aquarium-reared specimens of *Myaka myaka* and *Pungu maclareni* were dissected according to the ethical guidelines. Dissected retinae were flash-frozen in liquid nitrogen and then stored at -80 °C before use. Nuclei isolation was performed according to the 10x Genomics Demonstrated Protocol (doc. no. CG000366, rev. D, 10x Genomics 2022) with the following modifications: (i) in the final resuspension step of the nuclei isolation protocol, nuclei were resuspended in a resuspension buffer (1x PBS, 2% BSA, 0.2 U/μl RNase inhibitor) instead of Diluted Nuclei Buffer, (ii) a lower concentration of RNase inhibitor (0.1 U/μl) was used in all other buffers (Wash Buffer, Lysis Buffer, Lysis Dilution Buffer), and (iii) the total incubation time in the lysis step was 4 minutes. Isolated nuclei in the resuspension buffer were placed on ice and transported to the sequencing facility (30 min). Nuclei were then stained with Hoechst 33342 stain (final concentration 10 μg/ml) and propidium iodide (final concentration 5 μg/ml), and sorted on a BD Influx Cell Sorter into a new tube containing chilled resuspension buffer. Propidium iodide was excited with a 561 nm laser and detected using a 610/20 nm bandpass filter. Hoechst was excited with a 355 nm UV laser and detected using a 455/50 nm bandpass filter. The sorted fraction of nuclei was subsequently concentrated via centrifugation – the sample was spun down at the speed of 600 RCF for 5 minutes at 4 °C, excess supernatant was discarded, and nuclei were resuspended in the remaining resuspension buffer (the desired volume of resuspension buffer to be left in the tube was estimated based on the nuclei count during the sorting). Sample quality, as well as nuclei concentration and viability, was then assessed under the microscope. Nuclei were stained with trypan blue and counted across multiple visual fields. Nuclei concentration was calculated as the average number of nuclei per field divided by the field volume. For the sample of *Myaka myaka*, concentration was established as 600 nuclei/μl and viability as 85%. The sample of *Pungu maclareni* had a concentration of 1031 nuclei/μl and viability 90%. Both samples were processed with the 10x Genomics Chromium Next GEM Single Cell 3ʹ Reagent Kit v3.1 (Dual Index) according to the official 10x Genomics protocol (doc. no. CG000315, rev. A, 2020) using the 10× Genomics Chromium platform. *Myaka myaka* library was prepared with Chromium Controller and *Pungu maclareni* library with Chromium iX (both 10x Genomics). The volume of cell suspension stock for each reaction was calculated to recover the target of 6000 nuclei. Post-construction quality of both libraries was assessed with the Bioanalyzer. Both libraries were sequenced on the Illumina platform, *Myaka myaka* with Illumina NextSeq 500/550 High-Output v2.5 Kit (75 cycles) and *Pungu maclareni* with NextSeq 1000/2000 P2 XLEAP-SBS Reagent Kit (100 Cycles).

### Single-nucleus RNA-seq analyses

Single-nucleus RNA-seq reads from *Myaka myaka* and *Pungu maclareni* retinae samples were mapped to the curated *Myaka myaka* genome. Although mapping *P. maclareni* reads to the *M. myaka* genome could potentially introduce minor species-specific biases, the two species are closely related, and this genome assembly constitutes the highest-quality reference available. Using a common reference ensured consistent gene annotation and comparability across datasets. For the indexing and mapping steps we used SimpleAF [71]. This tool integrates Piscem, to generate a splice-aware index (i.e., spliceu) from the *Myaka* unmasked genome and annotations, and Alevin-Fry for efficient pseudoalignment and quantification using the following options: --chemistry 10xv3, --unfiltered-pl 10xv3_whitelist.txt, --min-reads 2000, --resolution parsimony-em, --use-piscem. The indexing step generated a default “transcript-to-gene” table, which we modified by replacing gene IDs with gene names (--t2g-map index/index/t2g_names.tsv), aiding subsequent analyses. The resulting count matrices were subsequently analyzed in Rstudio using the Seurat v.5 package [72] for quality control, normalization, dimensionality reduction, clustering, and visualization. In brief, matrices were imported using the fishpond package in snRNA mode and converted into Seurat v5 objects. Initial quality control involved filtering nuclei with fewer than 200 detected genes, more than 2,500 genes, a mitochondrial gene content above 3%, or abnormal transcript complexity (log₁₀ genes per UMI ≤ 0.9). Filtered datasets were normalized, variable features identified, and scaled while regressing out mitochondrial content. Dimensionality reduction was performed using PCA, and cell clustering was conducted with FindNeighbors and FindClusters; UMAP was used for visualization. Ambient RNA contamination was identified and corrected using the scCDC framework [73], and putative doublets were detected and removed using DoubletFinder [74] with empirically optimized parameters. The *P. maclareni* dataset exhibited slightly cleaner profiles compared to *M. myaka*, showing lower ambient RNA contamination ratio (0.0056 vs. 0.0059) and fewer detected doublets (398 vs. 653). Corrected singlet datasets were re-normalized using SCTransform, followed by PCA, clustering, and UMAP. Differential gene expression (DGE) analysis across clusters was performed with FindMarkers, retaining genes expressed in at least 25% of cells with a log fold-change >0.25 and adjusted p-value <0.05, and visualized using DotPlot. These DGE results were used for cluster annotations, assisted by an additional check of known retinal marker genes in Zebrafish that were curated from the literature [29]. This approach facilitated cell type characterization and comparison across species. The *P. maclareni* dataset included exclusively retinal cell identities. In contrast, *M. myaka* also yielded a small fraction of non-retinal cell types (e.g., vascular endothelial cells), likely reflecting deeper dissection. The final filtered and corrected Seurat objects for both *M. myaka* and *P. maclareni* contained good quality singlet retinal nuclei suitable for downstream analyses. Both datasets were integrated using SCTransform normalization. Integration features were selected, objects prepared with PrepSCTIntegration, and anchors identified to align datasets with IntegrateData. The integrated data was then analyzed with PCA, UMAP, and clustering to visualize shared and species-specific cell populations. We followed the same strategy for annotating cell types as previously described, performing a DGE analysis and also checking the list of known retinal markers as additional evidence. This approach allowed us to identify cell types for both species. Since this study is focused on cone cells, we subsetted the corresponding cluster and performed cross-species and species-specific analysis in order to characterize different cone types (LWS, RH2, SWS) based on different transcriptional profiles. To render plots we used Seurat (i.e., DimPlot, FeaturePlot, and DotPlot) and ggplot2 in Rstudio.

To assess conserved photoreceptor identity, we compared expression of the conserved marker sets proposed by Tommasini et al. (2025) [19] across cone populations in *P. maclareni* and *M. myaka*. We included all marker genes present in the target genome, including annotated paralogs, and visualized their scaled mean expression across cone populations using heatmaps, with genes ordered according to the PR1–PR4 categories defined by Baden et al. (2025) [10]. To characterize transcriptional differences between species, we additionally compared *P. maclareni* LWS versus *M. myaka* RH2, *P. maclareni* RH2 versus *M. myaka* RH2, and *P. maclareni* LWS versus *P. maclareni* RH2 populations. For each gene, we calculated mean expression, the percentage of cells expressing the gene, and a descriptive log2 fold-change. Because each species was represented by a single biological sample, these comparisons were treated as exploratory/descriptive rather than formal differential-expression analyses, and no statistical significance or FDR was calculated.

We further characterized the relative expression of the two *RH2A* paralogs, *opn1mw4-3* (*RH2Aα*) and *opn1mw4-2* (*RH2Aβ*), by classifying individual cones according to their relative paralog expression. For each cone cell, normalized RNA expression values were extracted and a paralog dominance score was calculated as D=(α−β)/(α+β), where positive and negative values indicate relative *RH2Aα* and *RH2Aβ* dominance, respectively. Cells with D > 0.2 were classified as α-dominant, cells with D < -0.2 as β-dominant, and cells with -0.2 ≤ D ≤ 0.2 as approximately balanced. These categories were used to quantify the distribution of relative *RH2A* paralog expression across cone cells. The top 50 cells at each extreme were used for downstream DGE analyses to characterize transcriptional differences associated with *RH2A* paralog dominance. For differentially expressed genes lacking gene names in our annotation file, we searched for corresponding entries in the NCBI Gene database. To verify that the two *RH2A* paralogs could be mapped unambiguously, we visually inspected their sequences, including the 3’UTR region, where most of the reads mapped. The paralogs showed sufficient sequence divergence in this region, and 93% of reads mapping to either paralog overlapped the UTR. Visual inspection of the mapped reads revealed no evidence of misassignment between paralogs (Fig. S4).

### Comparative regulatory analysis of the *tbx2a* locus

To investigate transcriptional regulation underlying differential *tbx2a* expression, we applied the pySCENIC [75] workflow to a combined single-nucleus RNA-seq dataset including *M. myaka* RH2 cones and *P. maclareni* LWS and RH2 cones. Cells were annotated by species and cone subtype, and highly variable genes were identified, with *tbx2a-1* retained in the expression matrix. A curated list of retinal transcription factors derived from zebrafish (https://www.ezrc.kit.edu/tfdb/go_query.php?go=retina) was filtered to retain only genes present in the dataset (n=38). Gene regulatory networks were inferred using GRNBoost2 based on co-expression patterns, followed by motif enrichment analysis using RcisTarget with mouse (mm10) ranking databases due to the lack of species-specific resources. To ensure compatibility with the motif database, gene names were standardized to mouse orthologs and paralogs were collapsed prior to analysis. Regulon activity was quantified at the single-cell level using AUCell, and aggregated by species and cone type to compute mean activity and the fraction of active cells. Differences in regulon activity between populations were assessed using Kruskal-Wallis tests followed by pairwise Wilcoxon rank-sum comparisons with Benjamini-Hochberg correction. Due to limited overlap between cichlid gene models and the reference motif database, only a subset of regulons was retained, and results were interpreted conservatively. To investigate regulatory variation at the *tbx2a-1* locus, we performed a comparative analysis focused on genomes of cichlid species, including *Metriaclima zebra* (Mzebra_GT3a), *Pungu maclareni* (ASM4175732v1), and *Myaka myaka* (this study). We performed BLASTN searches to identify *tbx2a* orthologs using *M. myaka tbx2a-1* sequence as a reference. The top hits across distinct scaffolds were retained and manually inspected in a genome browser (IGV). Orthology assignment of *tbx2a-1* in *M. zebra* was confirmed using known syntenic markers *acaca* and *pdcl1*. For each *tbx2a-1* locus, genomic regions spanning 20 kb upstream and 5 kb downstream of the gene were extracted using SAMtools. Gene sequences and flanking regions were then aligned using MAFFT (v7.520), and alignments were visually inspected in Geneious Prime®. Based on these alignments, a region approximately 9 kb upstream of the transcription start site was selected for detailed analysis, as it contained lineage-specific insertions in *M. myaka* not present in the other species. To identify putative regulatory differences, transcription factor binding sites (TFBS) were predicted using FIMO [76] with the JASPAR 2024 vertebrate motif database [77], using a significance threshold of 1 × 10⁻⁴. Motif scanning was performed on strand-corrected upstream sequences. Predicted TFBS were compared across species and projected onto the multiple sequence alignment to identify conserved and lineage-specific motifs, with particular emphasis on motifs unique to *M. myaka*. For visualization in Fig. 2C, a curated subset of biologically relevant TFs was selected based on prior knowledge in zebrafish: cone identity TFs (THRB, RXRG, MAFB, SIX7, GDF6A, TBX2), *tbx2a* upstream TFs (PITX2, FOXC1, FOXC2, GATA4, GATA5, GATA6, TBX3, TBX5, T, SMAD2, SMAD3, SMAD4, GLI2, GLI3, MEIS1, PBX1, HAND2), vision/opsin TFs (CRX, OTX2, OTX5, RORB, RORA, THRB, SIX3/6, SIX7, RAX, RAXL1, NR2E3, NRL, VSX1, VSX2, ESRRG), repressors (REST, ZEB1, ZEB2, HES1, HEY1, HEY2, TCF7L2, BHLHE40, BHLHE41, KLF9, KLF10) and developmental TFs (SOX2, POU4F1, POU4F2, FOXN4, ASCL1, ATOH7, NEUROD1). In the figure, only TFBS from this filtered subset are shown as black horizontal lines to improve readability, while the accompanying histograms quantify the total number of all predicted TFBS per species within each focal region. All predicted TFBS are listed in Table S5. Following the same approach, we scanned the upstream regions of the *LWS*, *RH2Aα*, and *RH2Aβ* opsins in *Myaka myaka* to identify potential TBX2-family binding sites and record their presence or absence.

### Whole-mount fluorescent in situ hybridization (FISH)

Whole-mount fluorescent in situ hybridization (FISH) was performed following Dalton et al. (2017) [3] to detect the opsin genes *RH2Aα* and *RH2Aβ*. The lower-expressed *RH2Aβ* was labeled with fluorescein (FL), and *RH2Aα* with digoxigenin (DIG). Total RNA was isolated from the retina of *M. myaka*, and double-stranded cDNA was synthesized using the Mint-2 cDNA synthesis kit (Evrogen). Gene-specific primers containing the T3 RNA polymerase promoter were used to amplify probe templates. Excess primers and other PCR by-products were removed by gel electrophoresis, and the purified fragments were recovered with the Gel/PCR DNA Fragments Extraction kit (Geneaid). RNA probes were transcribed in vitro using T3 RNA polymerase, followed by DNase I treatment to remove template DNA, and EDTA to stop the reaction. Retinas were dissected, fixed overnight in 4% PFA, transferred, and stored in 100% methanol until use. Retinas were cleared in xylene, rehydrated, and digested with Proteinase K before postfixation in 4% PFA. Samples were subsequently transferred into a hybridization solution for 2 hours and then incubated overnight at 56°C with FL- and DIG-labelled riboprobes. After hybridization, retinas were washed several times in SSCT at 65 °C and blocked for 2 hours in 1X Roche Molecular Blocking reagent. Detection was performed first with horseradish peroxidase (HRP)-conjugated antiFL antibody (overnight at 4°C) followed by washes in maleate buffer and signal amplification using the Tyramide SuperBoost™ Kits with Alexa Fluor™ 488 Tyramides (ThermoFisher Scientific). The procedure was repeated with anti-DIG antibodies and signal amplification with Tyramide SuperBoost™ Kits with Alexa Fluor™ 594 Tyramides (ThermoFisher Scientific). Finally, retinas were cleared in 70% glycerol and mounted on slides. Images were acquired on aZeiss LSM 880 scanning confocal microscope at the Viničná Microscopy Core Facility in Prague.

### Seasonal variation in opsin gene expression

Adult *Myaka myaka* specimens were collected as described above during both dry and rainy seasons. A total of 14 retina transcriptomes were generated (Table S1). Following dissection, retinal tissue was preserved in RNAlater™, stored at room temperature during fieldwork, transferred to a field refrigerator, and subsequently stored at −80 °C upon arrival in the laboratory. RNA was extracted from the dissected retinas using Qiagen RNeasy Micro Kit (www.qiagen.com). The quality and quantity of extracted RNA were verified using a 2100 Bioanalyzer (Agilent) and Qubit Fluorometer (Thermo Fisher Scientific) respectively. Libraries preparation and sequencing has been outsourced to Novogene Co., Ltd. Sequenced libraries were quality-checked using FastQC (http://www.bioinformatics.babraham.ac.uk/projects/fastqc) and relative opsin gene expression was calculated using Geneious v.9.1.4. For each sample, the pair-end library has been mapped against the seven cone opsin genes (*SWS1, SWS2B, SWS2A, RH2B, RH2Aβ*, *RH2Aα, LWS)* from *M. myaka*. The reference sequences were generated by mapping transcriptome reads to annotated Nile tilapia (*Oreochromis niloticus*) opsin gene sequences (*SWS1*, *opn1sw1*, LOC100710249; *SWS2B*, *opn1sw2*, LOC100695018; *SWS2A*, LOC100695287; *RH2B*, LOC100711209; *RH2Aβ*, LOC100710942; *RH2Aα*, LOC100710676; and *LWS*, LOC100694761). To differentiate *RH2Aβ* and *RH2Aα* paralogs, libraries were mapped to the fourth exon, where distinguishing SNPs are located. Based on the number of mapped reads to each reference gene the FPKM (fragments per kilobase and million reads) were calculated, also taking into account the library size and the length of each gene. Proportional opsin expression was calculated as a percentage of total relative expression for single-cone and double-cone classes separately. Differences in proportional expression between rainy- and dry-season *M. myaka* individuals were tested using beta-regression, allowing the use of non-transformed data in percentages and proportions, following Stieb et al. (2016) [78] and Kłodawska et al. (2025) [18]. Performed with the R package betareg v3.2-1 [79], this analysis is suitable for fitting non-transformed dependent variables within the (0,1) interval. The effect of season (i.e., depth) was tested for RH1 and selected cone opsin gene classes separately.

### Retinal wholemount preparation for photoreceptor and ganglion cells examination

Retinal wholemounts were prepared following standard protocols [47,80]. Eyes were dissected, and several radial cuts were made to allow the retina to lie flat on a slide. Orientation was confirmed using the dorsal cut and the position of the falciform process. The sclera, choroid, and retinal pigment epithelium (RPE) were gently removed using forceps and fine paintbrushes. To further eliminate residual RPE, retinas were bleached either in 3% hydrogen peroxide in PBS or, when manual removal was difficult, using a peroxide-formamide solution under a lamp (500µl formamide, 250µl 20x SSC, 2.8ml 30% peroxide, distilled water to 10ml). Samples were then rinsed three times for 5 min in PBS. For photoreceptor analysis, retinas were flat-mounted in glycerol on microscope slides, with the photoreceptors facing up. For ganglion cell analysis, retinas were mounted photoreceptor-side down and exposed to 37% formaldehyde vapor overnight at room temperature to enhance Nissl staining[80]. Wholemounts were then rehydrated, stained for 3 min in 0.1% cresyl violet, dehydrated in an ethanol series, cleared in xylene, and mounted in Entellan New (Merck).

### Stereological analysis and topographic map reconstruction

The topographic distribution of the photoreceptors was assessed using the optical fractionator technique[81], as adapted for retinal wholemounts by Coimbra et al. (2009) [82]. Each retina was treated as a single section; thus the thickness sampling fraction (tsf) was fixed at 10µm. The retinal outline was traced in Stereo Investigator (Microbrightfield, USA) using a 10x objective on an Olympus BX51 microscope equipped with a motorized stage and a digital camera. Photoreceptors were then systematically and randomly counted with a 100x oil-immersion objective according to the parameters listed in Table S8.

Photoreceptor counting proved challenging due to the difficulty of removing the RPE without damaging underlying layers of the retina, and because even after bleaching the photoreceptor layer often remained insufficiently clear for consistent quantification. Consequently, only four Barombi Mbo species, with three individuals per species, were included in this analysis. To ensure comparable sampling effort across individuals of different retinal sizes, the sampling grid size was adjusted to yield approximately 200 sites per retina, allowing Schaeffer coefficients of error (CE) below the accepted threshold of 0.1 [83,84].

Ganglion cells counting also presented some difficulties. Three types of cells can be identified based on cytological criteria from Nissl stained samples: ganglion cells, amacrine cells, and glial cells[85,86]. In the studied species, it was sometimes difficult to confidently distinguish between amacrine and the ganglion cells; therefore both cell types were counted. This approach does not affect the reconstruction of topographic maps, as previous studies have shown that high-density areas of amacrine and ganglion cells largely overlap [80,85,87]. Including amacrine cells in ganglion cell counts may result in a slight overestimation of total cell density. However, because amacrine cells were present at relatively low densities, this effect is minimal. In regions of high retinal cell density, additional sub-sampling was performed by halving the grid size while keeping the counting frame unchanged. This approach allowed for a more precise estimation of peak ganglion cell density. This part of the study included eight Barombi Mbo cichlid species, with two to three individuals per species. Cell counting was performed following the same methodology as described for photoreceptor analysis, using the parameters listed in Table S8.

Topographic density maps were constructed using R Studio software (R Foundation for Statistical Computing 2012) using data exported from the Stereo Investigator v.11.06.2 (32-bit) (https://www.mbfbioscience.com/products/neurolucida) according to Garza-Gisholt et al. (2014) [88]. The Gaussian kernel smoother from the Spatstat package was used[89] and the sigma value was adjusted to the grid size.

### Calculation of spatial resolving power from ganglion cell density

The upper limit of spatial resolving power (SRP) for each species was determined by using the peak density of ganglion cells [52]. To calculate the angle α, which represents the angle subtended by 1 mm on the retina, we used Matthiessen’s established ratio, stating that the focal length (f) in teleost fishes is approximately 2.55 times the radius of the lens [90,91]:

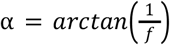

With the peak ganglion cell density (PGC, in cells/mm²), and angle α, we estimated the spatial resolving power (SPR) in cycles per degree (cpd). Since at least two ganglion cells are required to distinguish the boundary of a black-and-white cycle of a grating, the SPR was calculated as:

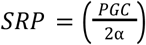

This provides an estimate of the upper limit of spatial resolving power.

### Histology and Transmission Electron Microscopy (TEM)

Eyes were dissected to isolate retinal tissue from five regions: central, dorsal, temporal, ventral, and nasal. The samples were then placed in PBS and sent to the Laboratory of Electron Microscopy, Faculty of Science, Charles University (Prague), for processing and resin embedding. Semi-thin sections were first examined, and subsequently, selected samples were mounted on copper grids for imaging with a Transmission Electron Microscope (TEM) for qualitative purposes.

## Acknowledgements

We are grateful to the Ministry of Scientific Research and Innovation in Cameroon to grant us research permits cited above and we thank the local Barombi village community for their permission to conduct our research on their land. We thank Antoine Pariselle and Ngando Ephesians Jiku for fieldwork assistance and Cyrille Dening for logistical support. We thank Dan Elleder for the high-molecular DNA extraction. We thank Michal Kolář and his team for assistance and expertise regarding the single-nucleus RNA sequencing. We thank Kristýna Eliášová for laboratory support. We thank Emilia Santos for her suggestions regarding the regulatory network analyses. We thank Veronika Truhlářová for providing the fish illustrations included in the figures.

This work was financially supported by the Grant Agency of Charles University (GAUK) 1524119 “Retinal specializations and visual adaptations with the link to the ecology of two teleost fish families: cichlids and mormyrids” (M.K., Z.M.), and Primus (Z.M.), the Fisheries Society of the British Isles (FSBI) travel grant (M.K.), the Czech Science Foundation grant 21-31712S “Zoom in the fish eye & blood: molecular evolution of functional adaptations in deep-sea and freshwater fishes” (Z.M.), the European Research Council (ERC) Consolidator grant (no. 101122542) “SensingDEEP” (Z.M.), the institutional support for the long-term development plan of the Military Health Institute of the Czech Republic (P. P.), the Australian Research Council with the Discovery Early Career Researcher Award DE200100620 and the LIEF grant LE100100074 supporting the QBI Advanced Microscopy Facility using Stereo Investigator (F.d.B). Computational resources were provided by e-INFRA CZ project 90254, supported by the Ministry of Education, Youth and Sports of the Czech Republic. Microscope imaging was co-financed by the Czech-BioImaging large research infrastructure project LM2023050.

## Data availability

Single-nuclei RNA-seq data, genome data, and bulk RNA-seq datasets have been deposited in the NCBI BioProject PRJNA1399146. All custom scripts used in this study are available in GitHub: https://github.com/FishEvoLab/Barombi_Mbo_single_cell.

## Author Contributions

M.K., P.B.G., and Z.M. conceived the study. M.K., A.I., A.R.B.N. and Z.M. collected samples. M.K. and Z.M. processed samples for bulk RNA-seq. P.B.G. and Z.K. processed samples for single-nucleus RNA-seq. P.P., D.E. and O.B. performed the genome sequencing and assembly. P.B.G. performed the genome annotation and single-nucleus RNA-seq analyses. D.B. assisted during the genome annotation. A.N. performed the FISH experiments. H.K. optimized the stereological methods. M.K. and F.d.B. performed the stereological analyses and generated topographic maps. M.K., P.B.G, and Z.M. drafted the manuscript, figures, tables, and SI Appendix. Z.M. supervised this study. All the coauthors contributed to the final version of the manuscript.

## Supplementary Information for

**This PDF file includes:**

Figures S1 to S5

Table S1 to S8 captions

## Figures S1 to S5

**Figure S1.**
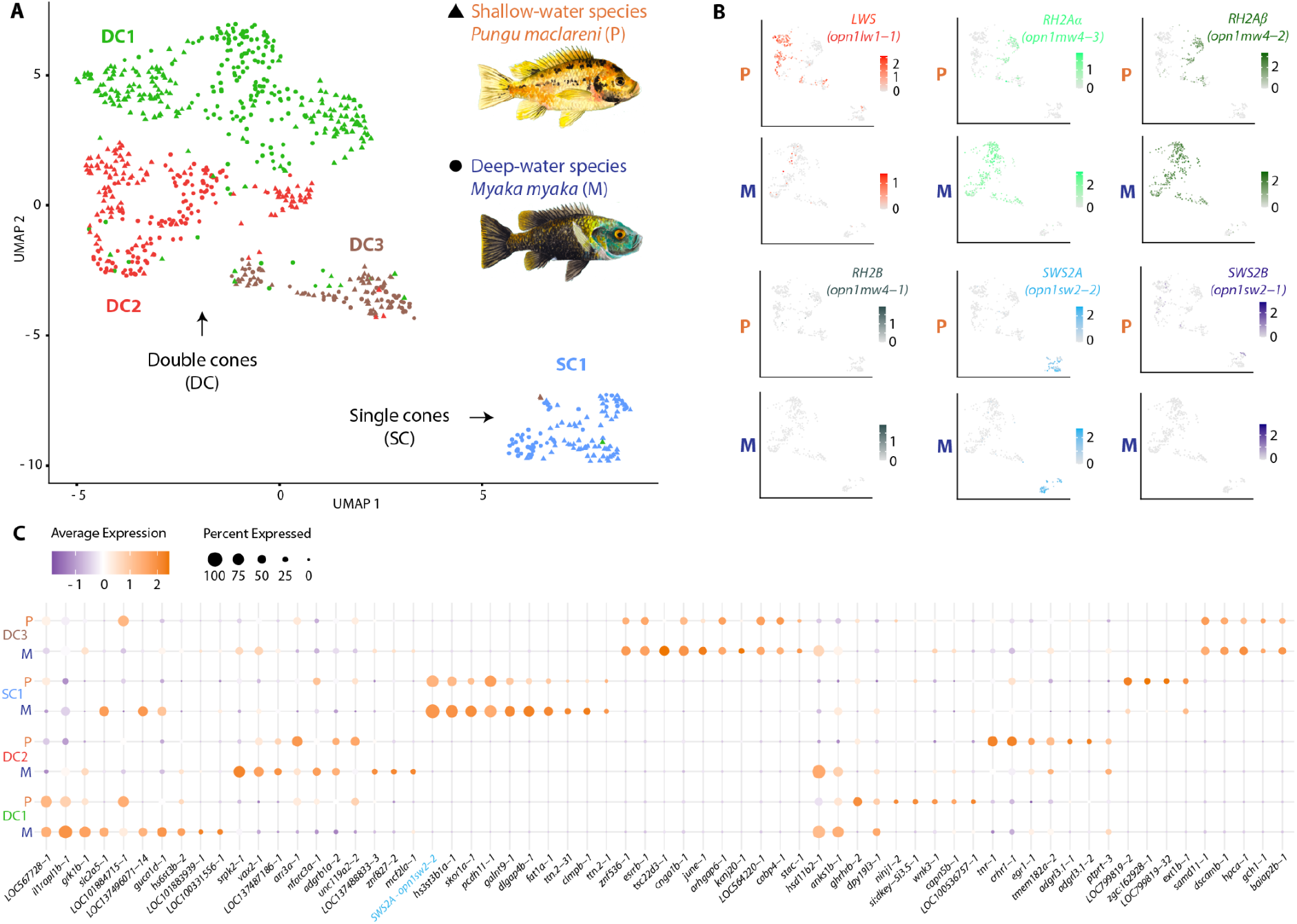
Transcriptional profiles of cone photoreceptors in deep- and shallow-water cichlids. **(A)** UMAP embedding of cone photoreceptor nuclei from *Myaka myaka* (deep-water species, dots) and *Pungu maclareni* (shallow-water species, triangles) integrated into a shared analysis, showing major cone clusters. **(B)** Feature plots showing normalized expression of opsin genes *opn1lw1-1* (*LWS*), *opn1mw4-3* (*RH2Aα*), *opn1mw4-2* (*RH2Aβ*), *opn1mw4-1* (*RH2B*), *opn1sw2-2* (*SWS2A*) and *opn1sw2-1* (*SWS2B*), in *M. myaka* (M) and *P. maclareni* (P). **(C)** Dot plot of the top 10 differentially expressed genes across cone clusters between species (table S4); color indicates average normalized expression and dot size represents the percentage of cells expressing each gene.

**Figure S2.**
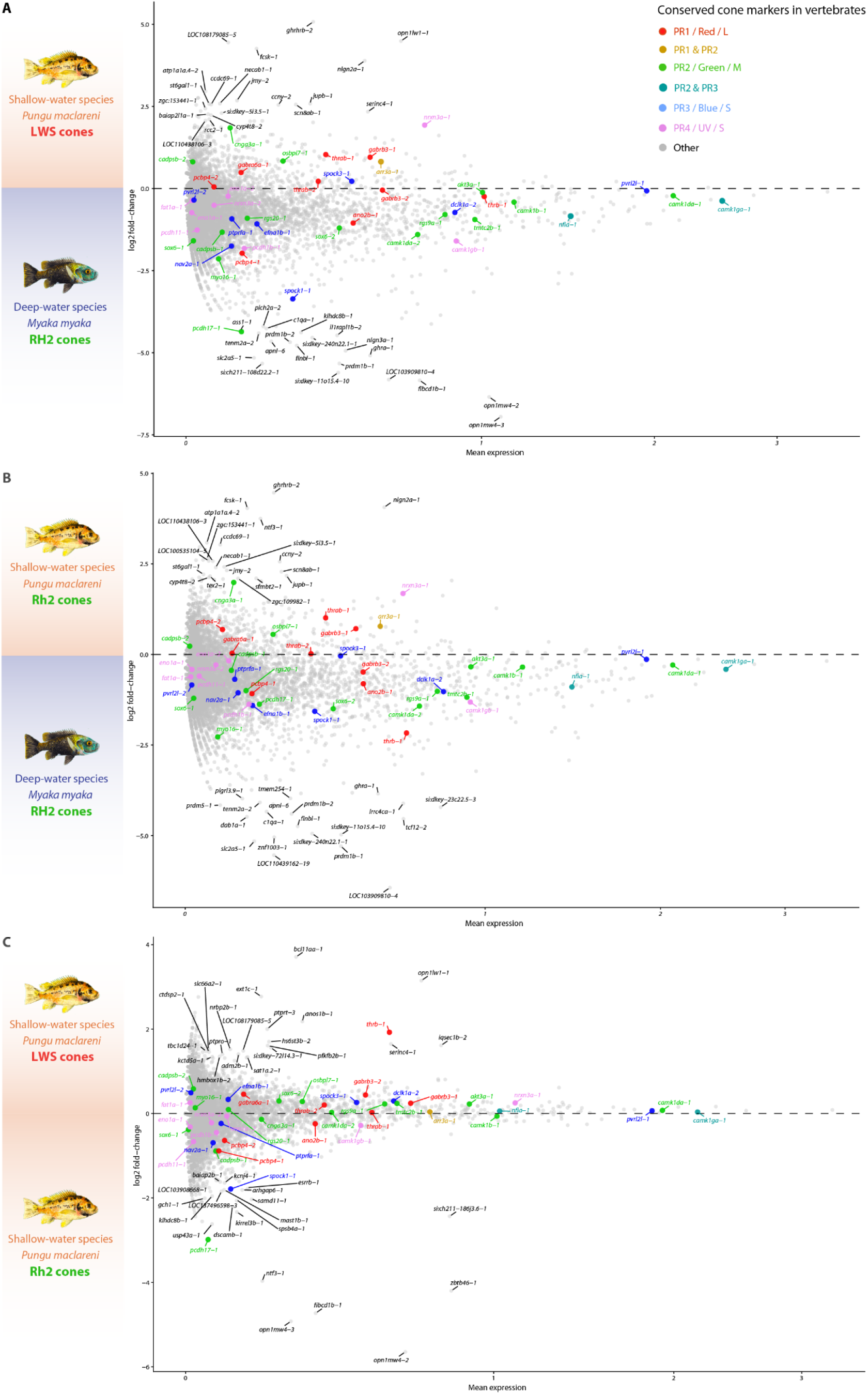
Comparison of conserved vertebrate cone photoreceptor markers between *Pungu maclareni* and *Myaka myaka*. Descriptive comparison of cone markers identified from Tommasini et al. (2025) [19] was examined between **(A)** *P. maclareni* LWS and *M. myaka* RH2 cells, **(B)** *P. maclareni* RH2 and *M. myaka* RH2 cells, and **(C)** *P. maclareni* LWS and *P. maclareni* RH2 cones. Points show mean normalized expression and the y-axis represents the descriptive log2 fold-change between populations. Conserved vertebrate cone markers are highlighted according to their assigned photoreceptor programme, and the top 20 other genes ranked by absolute log2 fold-change are labelled for the cone population of each species.

**Figure S3.**
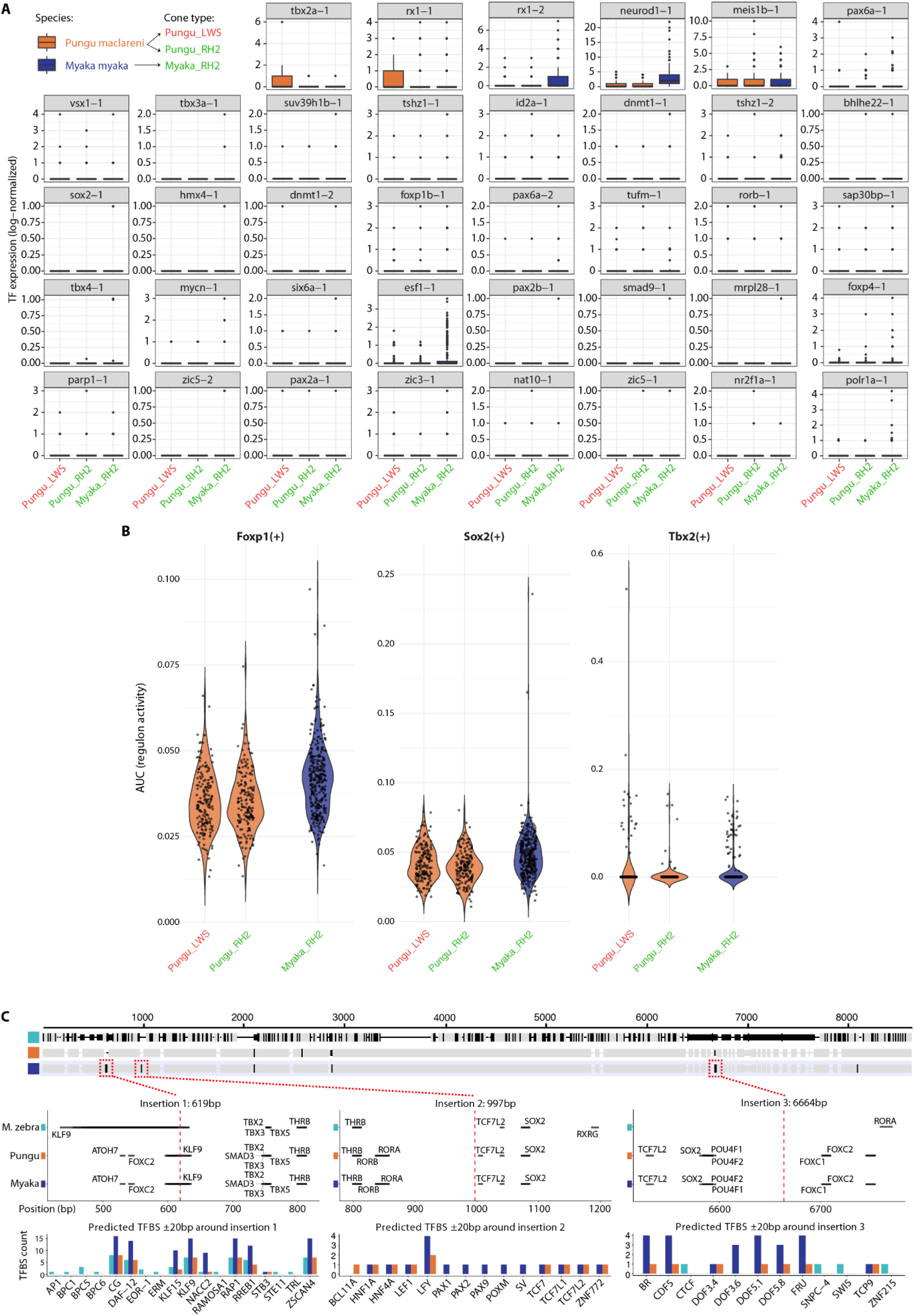
Regulatory and cis-regulatory landscape of *tbx2a* in deep-water cichlids. **(A)** Expression of transcription factors previously identified in zebrafish retina across double-cone populations (LWS and RH2) in *P. maclareni* and *M. myaka*, shown as boxplots. **(B)** pySCENIC-inferred transcription factor activity across LWS and RH2 cone populations, shown as violin plots for *Foxp1*, *Sox2*, and *Tbx2*. Activity is measured as Area Under the Curve (AUC), reflecting coordinated expression of predicted target genes in individual cells (higher values indicate higher inferred activity). *Tbx2* activity is slightly reduced in RH2 cones relative to LWS cones, whereas *Foxp1* and *Sox2* remain stable across species and cone types, suggesting locus-specific modulation of *Tbx2* regulatory activity rather than global transcriptional shifts. **(C)** Motif enrichment in upstream regulatory regions of *tbx2a* across *Metriaclima zebra Pungu maclareni*, and *Myaka myaka*. Sequences were aligned to enable positional comparison across species. For readability, the figure shows transcription factor binding sites (TFBS) from a filtered set of biologically relevant transcription factors (TFs) (see Methods); all predicted TFBS are listed in table S5. Each black line indicates a predicted TFBS mapped onto the alignment (x-axis: alignment position; y-axis: species). Vertical dashed red lines indicate the positions of lineage-specific insertions identified in *M. myaka.* The proximal insertion (+16 bp at –612 bp) is associated with local enrichment of *klf9* binding sites, whereas more distal insertions (+7 bp at –970 bp; +14 bp at –6,000 bp) generate fewer relevant TFBS. Histograms summarize total predicted TFBS counts per species in each region.

**Figure S4.**
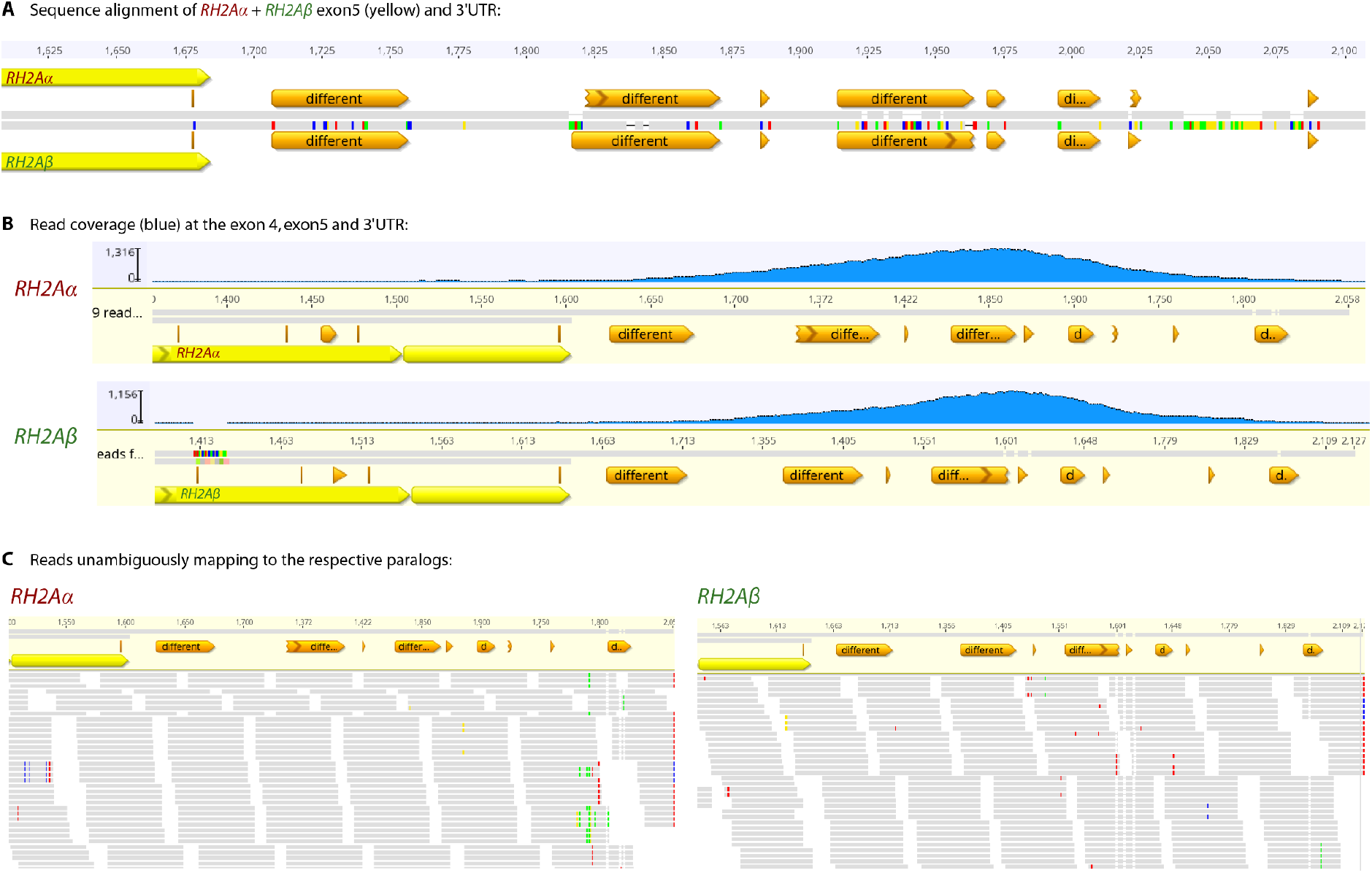
Control for unambiguous mapping to the two *RH2A* paralogs. **(A)** DNA sequences of the two *RH2A* paralogs, including the 3’ UTR, were aligned. **(B)** Most reads mapped to the 3’ UTR, where the two paralogs show sufficient sequence divergence for unambiguous mapping; 93% of reads mapping to either paralog overlapped the UTR region. **(C)** No reads showed evidence of misassignment between the two paralogs. Grey indicates sequence identity, whereas coloured bases indicate sequence differences. Yellow annotations indicate coding sequences (exon 5 or exons 4–5), and orange annotations indicate other regions of the sequence.

**Figure S5.**
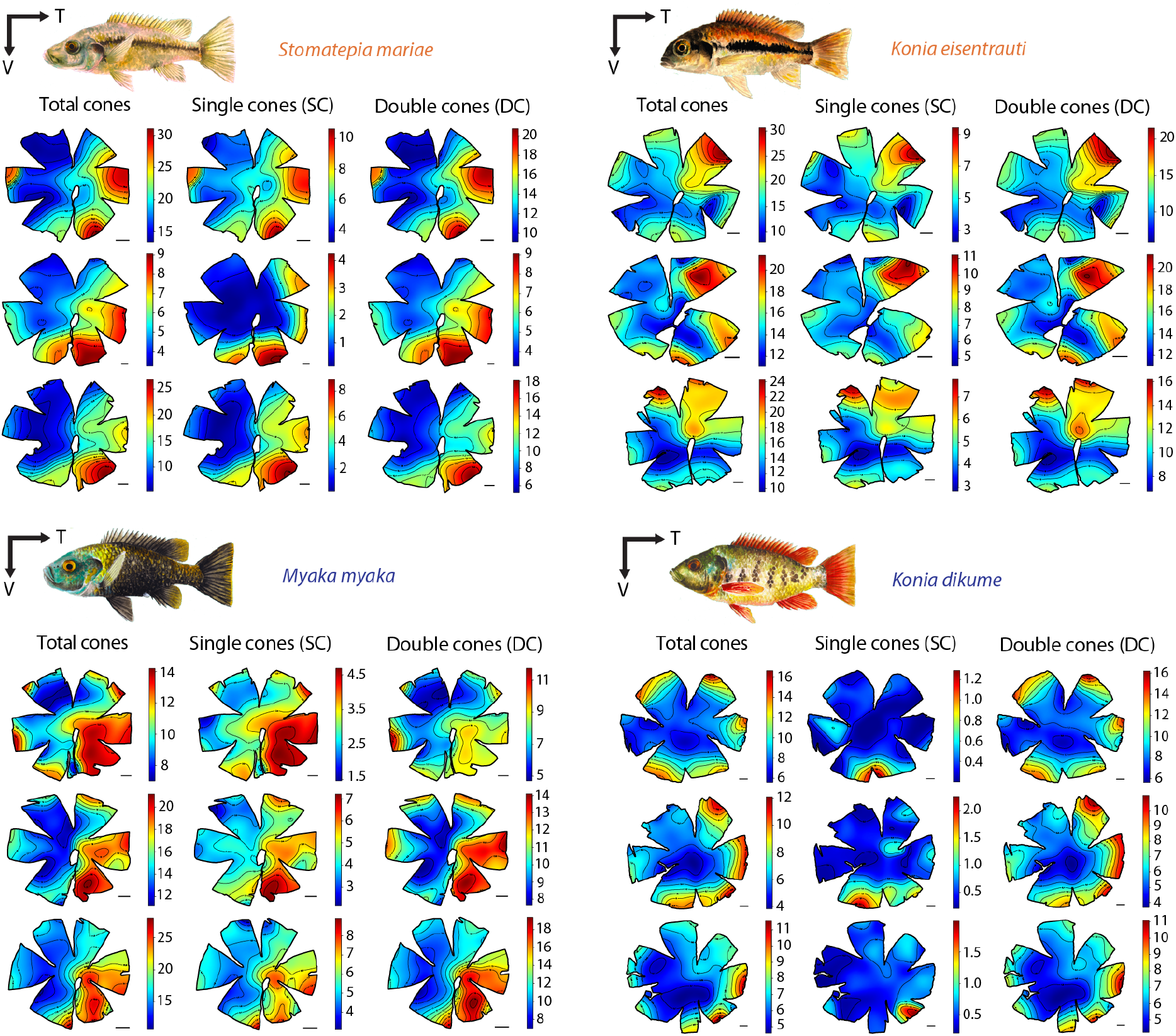
Retinal topographic maps of photoreceptor cells in Barombi Mbo cichlids. Photoreceptor cell density maps for shallow-water species (*Stomatepia mariae* and *Konia eisentrauti*) and deep-water species (*Myaka myaka* and *Konia dikume*). Three individuals per species are shown. T= temporal, V = ventral. Scale bars represent 1 µm.

## Table S1 to S8 captions

**Table S1.** Samples and datasets used in this study, including single-nucleus RNA-seq data, genome sequencing data, bulk RNA-seq data, and retinal stereological analyses quantifying photoreceptors and ganglion cells.

**Table S2.** Differentially expressed genes used for retinal cell cluster annotation. The table reports DGE results for each cluster, including gene name, P value, adjusted P value, average log₂ fold change, percentage of expressing cells (pct.1 and pct.2), cluster number, and assigned cell type.

**Table S3.** Marker genes of species-specific cone clusters in *Pungu maclareni* and *Myaka myaka.* The table lists differentially expressed marker genes identified in each species’ cone clusters, including statistics from the DGE analysis used for the dot plot in Fig. 2A and 2B.

**Table S4.** Differential gene expression analysis of cone clusters between *Pungu maclareni* and *Myaka myaka*. The table lists the top 10 differentially expressed genes per cone cluster, including statistics from the DGE analysis used for the dot plot in fig. S1C.

**Table S5.** Lineage-specific regulatory motifs in the 9 kb upstream regions of *tbx2a* in *Myaka myaka*, *Pungu maclareni*, and *Metriaclima zebra*. The table lists predicted transcription factor binding sites (TFBS) identified using FIMO (MEME Suite) in the 9 kb upstream region of the *tbx2a* locus in each species, highlighting potential cis-regulatory divergence.

**Table S6.** Differential gene expression in *RH2Aα*-versus *RH2Aβ*-dominant cones of *Myaka myaka* and *Pungu maclareni*. The table lists genes enriched in the 50 cells with the strongest relative expression of each *RH2A* paralog. Positive avg_log2FC values indicate genes enriched in *RH2Aα*-dominant cones, whereas negative values indicate genes enriched in *RH2Aβ*-dominant cones. Only the top 100 genes per species ranked by differential expression are listed for readability.

**Table S7.** Summary of the stereological parameters used for ganglion cell and photoreceptor topography analyses. Morphometric variables include lens diameter (W, mm), total length (TL, cm), and standard length (SL, cm). The precision of the stereological estimates is indicated by the Schaeffer coefficient of error (CE).

**Table S8.** Estimates of ganglion and photoreceptor cell abundance, including mean cell counts, variance measures, and sampling parameters derived using Stereoinvestigator. Metrics include estimated mean cell count, variance of cell counts, variance of the mean, contour area to counting frame area ratio, estimated variance of the total cell population, and coefficient of error (CE, Scheaffer). Finite population correction assumed to be 1¹. Estimate of maximum possible sampling sites².

